# Revealing unobserved factors underlying cortical activity using a rectified latent variable model applied to neural population recordings

**DOI:** 10.1101/072173

**Authors:** Matthew R. Whiteway, Daniel A. Butts

## Abstract

The activity of sensory cortical neurons is not only driven by external stimuli, but is also shaped by other sources of input to the cortex. Unlike external stimuli these other sources of input are challenging to experimentally control or even observe, and as a result contribute to variability of neuronal responses to sensory stimuli. However, such sources of input are likely not “noise”, and likely play an integral role in sensory cortex function. Here, we introduce the rectified latent variable model (RLVM) in order to identify these sources of input using simultaneously recorded cortical neuron populations. The RLVM is novel in that it employs non-negative (rectified) latent variables, and is able to be much less restrictive in the mathematical constraints on solutions due to the use an autoencoder neural network to initialize model parameters. We show the RLVM outperforms principal component analysis, factor analysis and independent component analysis across a variety of measures using simulated data. We then apply this model to the 2-photon imaging of hundreds of simultaneously recorded neurons in mouse primary somatosensory cortex during a tactile discrimination task. Across many experiments, the RLVM identifies latent variables related to both the tactile stimulation as well as non-stimulus aspects of the behavioral task, with a majority of activity explained by the latter. These results suggest that properly identifying such latent variables is necessary for a full understanding of sensory cortical function, and demonstrates novel methods for leveraging large population recordings to this end.

## INTRODUCTION

The sensory cortex not only represents information from the sensory periphery, but also incorporates input from other sources throughout the brain. In fact, a large fraction of neural activity in the awake sensory cortex cannot be explained by the presented stimulus, and has been related to a diversity of other factors such as stimulation of other sensory modalities (Ghazanfar and Schroeder, 2006; De Meo et al., 2015), location within the environment (Haggerty and Ji, 2015), and numerous aspects associated with “cortical state” (Harris and Thiele, 2011; Marguet and Harris, 2011; Pachitariu et al., 2015) including attention (Harris and Thiele, 2011; Rabinowitz et al., 2015), reward (Shuler, 2006) and state of arousal (Otazu et al., 2009; Niell and Stryker, 2010). Activity in sensory cortex linked to such non-sensory inputs can result in variability in the responses of neurons to identical stimulus presentations, which has been a subject of much recent study (Goris et al., 2014; Amarasingham et al., 2015; Rabinowitz et al., 2015; Cui et al., 2016). This suggests that a full understanding of sensory cortical function will require the ability to characterize non-sensory inputs to sensory cortex and how they modulate cortical processing.

However, such non-sensory inputs are typically not under direct experimental control nor directly observed, in which case their effects can only be inferred through their impact on observed neural activity. For example, shared but unobserved inputs can lead to noise correlations observable in simultaneously recorded neurons (Cohen and Kohn, 2011; Doiron et al., 2016), which can serve as a means to predict one neuron’s activity from that of other neurons (Schneidman et al., 2006; Pillow et al., 2008; Vidne et al., 2011). Noise correlations thus demonstrate one approach to understanding neural variability, and other recent extensions of this idea have used the summed activity of simultaneously recorded neurons (Okun et al., 2015; Schölvinck et al., 2015) and local field potentials (Rasch et al., 2008; Cui et al., 2016) to capture the effects of non-sensory inputs. Notably, these approaches all focus on the effects of shared variability on single neuron activity, and thus do not fully leverage the simultaneous recordings from multiple neurons to infer shared sources of input.

An alternative is to jointly characterize the effects of unobserved, non-sensory inputs on a population of simultaneously recorded neurons. This approach is embodied in a class of methods known as latent variable models (Cunningham and Yu, 2014), which aim to explain neural activity over the population of observed neurons using a small number of factors, or “latent variables”. Latent variable models evolved from classic dimensionality reduction techniques like Principal Component Analysis (PCA) (Ahrens et al., 2012; Kato et al., 2015), and encompass a wide range of methods such as Factor Analysis (FA) (Churchland et al., 2010), Independent Component Analysis (ICA) (Freeman et al., 2014), Poisson Principal Component Analysis (Pfau et al., 2013), Locally Linear Embedding (Stopfer et al., 2003), Restricted Boltzmann Machines (Koster et al., 2014), state space models (Smith and Brown, 2003; Paninski et al., 2009; Macke et al., 2011; Archer et al., 2014; Kulkarni and Paninski, 2015), and Gaussian Process Factor Analysis (Yu et al., 2009; Semedo et al., 2014; Lakshmanan et al., 2015).

Here, we propose a new latent variable approach called the Rectified Latent Variable Model (RLVM). This approach leverages two innovations over previous methods. First, it constrains the latent variables to be non-negative (rectified), which is hypothesized to be a fundamental nonlinear property of neural activity (McFarland et al., 2013) that can lead to important differences in the resulting descriptions of population activity (Lee and Seung, 1999). Indeed, using simulations, we show that rectification is necessary for the RLVM to recover the true activity of non-negative latent variables underlying population activity. The second innovation is that the RLVM avoids several statistical constraints on the latent variables that are necessary in other methods; for example, it does not require them to be uncorrelated (like PCA), independent (like ICA) or follow Gaussian distributions (like FA). To enable such unconstrained estimation of model parameters, we base solutions of the RLVM on an autoencoder (Bengio et al., 2013), which allows the RLVM to efficiently scale up to large datasets from both electrophysiological and optical recordings.

We first describe the RLVM and demonstrate its application it to a synthetic dataset generated to resemble typical large-scale recordings produced by 2-photon experiments. This synthetic dataset gives us “ground truth” with which to compare RLVM performance with a range of other latent variable approaches. We demonstrate that the RLVM outperforms these alternatives due to the innovations described above. We then apply the RLVM to a large 2-photon dataset recorded in mouse barrel cortex during a decision-making task (Peron et al., 2015). The relationship between the latent variables inferred by the RLVM and the behavioral observations related to the task revealed that a large proportion of cortical activity is related to non-vibrissal aspects of the behavioral task. Furthermore, consistent with the results on the synthetic dataset, the RLVM had the ability to match or outperform the other tested latent variable approaches, and also identified latent variables most correlated with individual observed aspects of the experiment. These results were consistent across many neural populations and animals sampled from this dataset, and thus identify consistent types of latent variables governing the diverse set of neurons recorded over many experiments. In total, this demonstrates that the RLVM is a useful tool for inferring latent variables in population recordings, and how it might be used in order to gain significant insights into how and why sensory cortex integrates sensory processing with non-sensory variables.

## METHODS

### Fitting the RLVM

The goal of the Rectified Latent Variable Model (RLVM) is to accurately predict observed neural activity **y**_*t*_ ∈ ℝ^*N*^ using a smaller set of latent variables 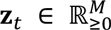. Here, **y**_*t*_ and **z**_*t*_ are vectors describing the activity at each time point *t*, and the matrices 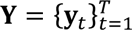 and 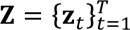 are all the observed data and latent variables, respectively, across time. The RLVM then tries to predict the observed activity **y**_*t*_ with the **z**_*t*_ as follows:

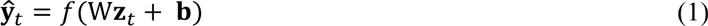

where *f*(.) is a parametric nonlinearity, and the model parameters are the coupling matrix W ∈ ℝ^*M*×*N*^ and the bias vector **b** ∈ ℝ^*M*^, collectively referred to as *θ* = {W, **b**}.

#### Estimation of model components

We estimate the model parameters *θ* and infer the latent variables **Z** using the maximum marginal likelihood (MML) algorithm, which is closely related to the Expectation-Maximization (EM) algorithm (Paninski et al., 2009; Vidne et al., 2011). The MML algorithm first infers the latent variables using initial model parameters 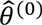, then updates the model parameters using the newly inferred latent variables. The algorithm continues to alternate between these two steps until a convergence criterion is met. Mathematically, each of these steps is defined as an optimization problem:

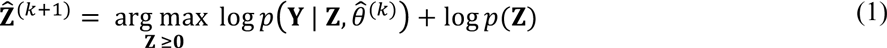

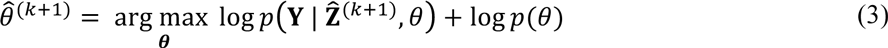

Both eqs. (2) and (3) correspond to a maximum a posteriori (MAP) estimate of the latent variables and model parameters, respectively, in which we maximize the sum of the data log-likelihood and a log-prior distribution. Although eq. (2) is a constrained optimization problem, it can be transformed into an unconstrained optimization problem as described below, and thus we solve both eqs. (2) and (3) using an unconstrained L-BFGS method.

The MML algorithm presented in eqs. (2) and (3) is a general procedure that can be specifically implemented for different types of data by properly defining the probability distribution in the data log-likelihood term log *p*(**Y** | **Z**, *θ*), which describes the probability of the observations given the current estimates of the latent variables and the model parameters. For example, in what follows we will use a Gaussian distribution for 2-photon data, but could instead use a Poisson distribution for spiking data. The forms of the log-prior terms log *p*(**Z**) and log *p*(*θ*) are in general independent of the form of the data log-likelihood term. Because this work is focused on the analysis of 2-photon data we’ll discuss the implementation of the MML algorithm that is specific to modeling 2-photon data, including a discussion of our treatment of the log-prior terms.

We first address the data log-likelihood terms of the form log *p*(**Y** | **Z**, *θ*). For 2-photon data, we model the observed fluorescence traces as a linear combination of the latent variables plus a bias term, 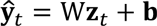 (eq. 1, with linear *f*(.)). Furthermore, we assume a Gaussian noise model so that 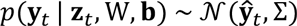 and

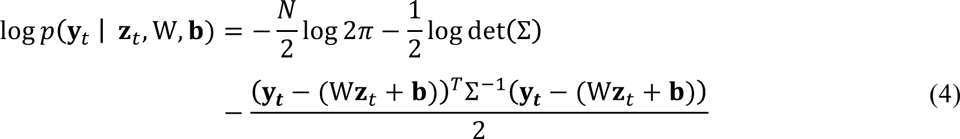

for a given time point *t*. For computational convenience we do not try to fit the noise covariance matrix ∑, but rather model it as a constant times the identity matrix. This constant can be incorporated into the log-prior terms, and hence does not show up in the final MML equations (eqs. 8 and 9 below). By modeling the noise covariance matrix as a multiple of the identity matrix we are making the assumption that the Gaussian noise has the same variance for each neuron (isotropic noise). Although not true in general, the advantage of this simplification is that we do not need to estimate the variance parameter, and (2) becomes a penalized least squares problem, which can be solved analytically. Constraining the noise covariance matrix to be diagonal (anisotropic noise) leads to solving a penalized *weighted* least squares problem, which must be solved iteratively.

We also make the assumption that data at different time points are conditionally independent, so that the full log-likelihood term can be written as

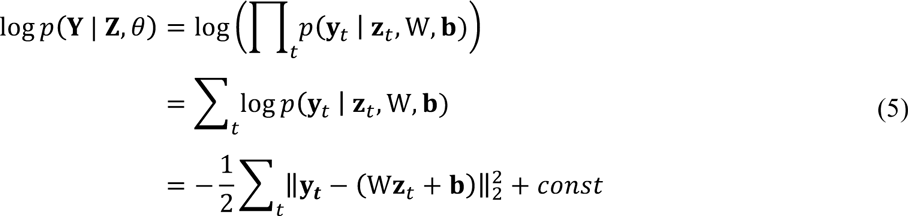

where 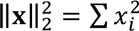 is the squared L_2_ norm of a vector x. The assumption of conditional independence is common practice when dealing with data log-likelihood terms (Bishop, 2006), and allows us to factorize the full conditional distribution log *p*(**Y** | **Z**, *θ*); without this assumption the resulting data log-likelihood term would be intractable.

To further constrain the types of solutions found by the model, we choose a particular form of the log-prior term log *p*(**Z**) (eq. 2). The are many different priors used for **Z** in the neuroscience literature on latent variable models, including latent dynamical systems priors (Paninski et al., 2009) and Gaussian Process priors (Yu et al., 2009). Here we use a simple smoothing prior that penalizes the second derivative of the time course of each latent variable **z**^*i*^, which can be written as

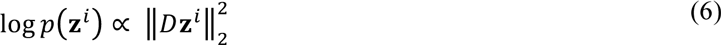

where *D* is the discrete Laplace operator. The full log-prior term log *p*(**Z**) is the sum of these terms for each individual latent variable.

The log-prior term log *p*(*θ*) in eq. (3) likewise allows for the incorporation of additional constraints on model parameters. We use a standard Gaussian prior on both the coupling matrix W and the biases **b**, so that

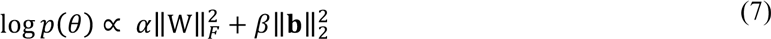

where 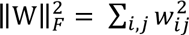 is the Frobenius norm of a matrix W and α and β are constants which scale the relative weight of each term. This prior has the effect of preventing the model parameters from growing too large, which can hurt model performance (Fig. A1).

Using the expressions in eqs. (5), (6) and (7), the 2-photon implementation of the general MML algorithm in eqs. (2) and (3) becomes

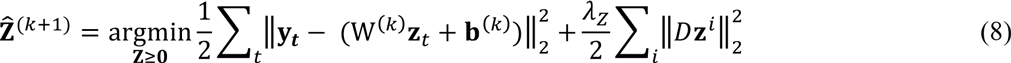

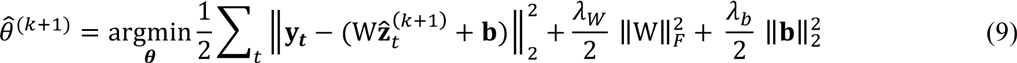

The λ values in front of the log-prior terms are hyperparameters that are chosen by hand (see the Model fitting section).

The non-negativity constraint on the latent variables **Z** is an important feature of the RLVM. Although it is possible to use explicitly constrained optimization techniques, we take a different approach that is more in line with the autoencoder optimization we use to obtain initial values for the MML algorithm (see below). Instead of optimizing non-negative latent variables **z**^*i*^, we substitute them with unconstrained latent variables **x**^*i*^ that are passed through a rectified linear (ReLU) function *g*(.):

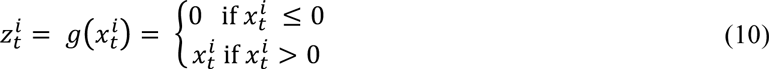

The model of neural activity (eq. 1) then becomes

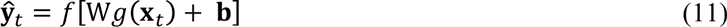

and unconstrained optimization techniques can be used in eq. (8) to solve for **X** instead of **Z**. Although the ReLU function is not differentiable at zero, we use the subdifferential approach common in the neural networks literature and define the derivative to be zero at zero (Hara et al., 2015).

#### Initialization of latent variables using an autoencoder

In the inference of **Z** (eq. 8), there are *T*M* parameters to estimate, which is a very high-dimensional space to search. The prior distribution we place on the latent variables is not a strong one, and as a result this optimization step tends to get stuck in poor local minima. In order to avoid these poor local minima, we start the MML optimization algorithm using initial estimates of both the model parameters and the latent variables from the solution of an autoencoder (Boulard and Kamp, 1989; Japkowicz et al., 2000; Bengio et al., 2013). An autoencoder is a neural network model that attempts to reconstruct its input using a smaller number of dimensions, and its mathematical formulation is similar to the RLVM; so similar, in fact, that the model parameters in the RLVM have direct analogues in the autoencoder, as shown below. Furthermore, the optimization routine for the autoencoder is faster and better-behaved than the MML algorithm, which makes it an attractive model for finding initial RLVM values.

The autoencoder takes the vector of neural activities **y**_*t*_ ∈ ℝ^*N*^ and projects it down onto a lower dimensional space ℝ^*M*^ using an *encoding matrix* W_1_ ∈ ℝ^*M*×*N*^. A bias term **b**_1_ ∈ ℝ^*M*^ is added to this projected vector, so that the resulting vector **x**_*t*_ ∈ ℝ^*M*^ is given by **x**_*t*_ = W_1_y_*t*_ + **b**_1_. W_1_ is said to *encode* the original vector **y**_*t*_ in the lower dimensional space with the vector **x**_*t*_, and this vector is analogous to the unconstrained latent variables in the RLVM. As with the RLVM, we enforce the non-negativity constraint on **x**_*t*_ by applying the ReLU function:

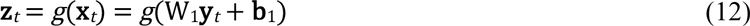

The autoencoder (again like the RLVM) then reconstructs the original activity y_t_ by applying a *decoding matrix* W_2_ ∈ ℝ^*N*×*M*^ to **z**_*t*_ and adding a bias term **b**_2_ ∈ ℝ^*N*^. The result is passed through a parametric nonlinearity *f*(.) so that the reconstructed activity 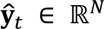 is given by

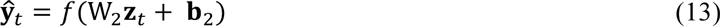

which matches the RLVM model structure in eq. (1). The weight matrices and bias terms, grouped as Θ = {W_1_, W_2_, **b**_1_, b_2_}, are simultaneously fit by minimizing the reconstruction error 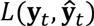 between the observed activity **y**_*t*_ and the predicted activity 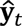:

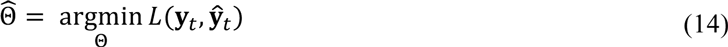

Once this optimization problem has been solved using standard gradient descent methods, we initialize the RLVM model parameters in eq. (2) with 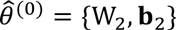. A notable feature of the autoencoder is that there is no need to alternate between inferring latent variables and estimating model parameters, as in eqs. (2) and (3); once the model parameters have been estimated using eq. (14), the latent variables can be explicitly calculated using eq. (12).

For modeling 2-photon data (as above), the noise distribution is Gaussian and the nonlinearity *f*(.) in eq. (13) is assumed to be linear. The reconstruction error 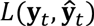 for Gaussian noise is the mean square error (again assuming equal noise variances across neurons), so in this special case of eq. (14) the autoencoder estimates for the weights and biases are given by:

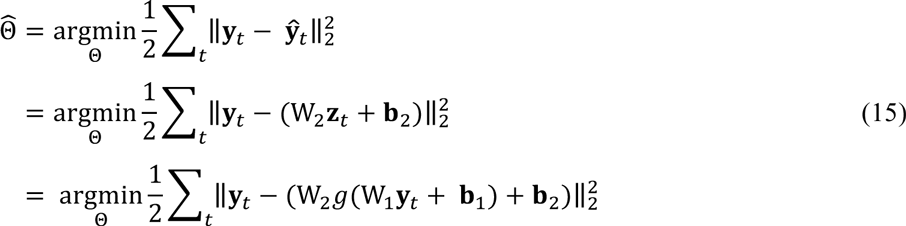

and we perform this optimization using an L-BFGS routine to obtain the weights and biases.

We also include regularization terms for the model parameters, which prevent over-fitting to the training data and can improve the model’s ability to generalize to new data (Bishop, 2006). As we saw previously, these regularization terms can also be interpreted as log-prior distributions on the model parameters in the probabilistic setting. A more general optimization problem for the autoencoder that includes both the reconstruction error and these regularization terms is

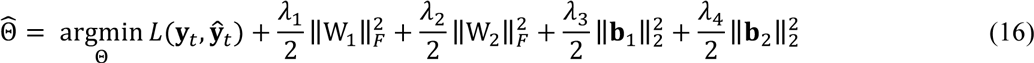

Large values of λ_*i*_ will encourage small values in the corresponding set of parameters. Furthermore, the use of regularization on the weight matrices helps to break a degeneracy in the autoencoder: because the reconstructed activity 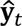 involves the product between the weights W_2_ and the latent variables z_t_ (eq. 13), an equivalent solution is given by the product of *c*^*^W_2_ and (1/c)^*^**z**_*t*_ for any positive constant c. Applying a regularization penalty to the weights W_2_ limits the range of values W_2_ can take and thus helps to set a scale for both the weights and the latent variables.

We also use “weight-tying” (Bengio et al., 2013), where the encoding and decoding weight matrices are constrained to be transposes of each other, i.e. W_2_ = (W_1_)^T^. This has the effect of nearly halving the number of model parameters that need to be estimated, which speeds up the model fitting procedure (Fig. A2). Not enforcing this constraint commonly results in qualitatively similar solutions (Fig. A3), and as a result, all models in this paper initialized with the autoencoder employ weight-tying.

### Model fitting

Model fitting using the MML algorithm requires alternating between inferring latent variables and estimating model parameters. We monitored the log-likelihood values throughout this procedure and ensured that the algorithm stopped only after additional iterations brought no further improvement. We compared the fitting behavior of the MML using random initializations versus autoencoder initializations (see Results). For these tests, we used the same regularization values for the latent variables (λ_*z*_ = 1) and for the model parameters (see below) to facilitate model comparisons.

The latent variable models we used to analyze the simulated and experimental data were the RLVM (regularization parameters set as λ_1_ = λ_2_ = 1000/(number of latent variables), λ_3_ = λ_4_ = 100, lambdas numbered as in eq. 16; code available for download at www.neurotheory.umd.edu/code), PCA (using MATLAB’s built-in function `pca`), FA (using MATLAB’s built-in function `factoran`; default settings), and ICA (using FastICA, available for download at http://research.ics.aalto.fi/ica/fastica/; default settings). The FA results reported using the dataset from Peron et al. (2015) used a PCA-based algorithm (available for download at http://www.mathworks.com/matlabcentral/fileexchange/14115-fa) rather than the maximum likelihood-based algorithm `factoran`, which proved prohibitively inefficient on such a large dataset. Autoencoder fitting was performed using a MATLAB implementation of the L-BFGS routine by Mark Schmidt, available for download at www.cs.ubc.ca/~schmidtm/Software/minFunc.html.

Model fitting was performed using 5-fold cross-validation (data is divided into five equally-sized blocks, with four used for training and one for testing, with five distinct testing blocks) unless otherwise noted. To assess the quality of model fits, we employed a procedure that is similar to the leave-one-out prediction error introduced in Byron et al. (2009). For all models (RLVM, PCA, FA, ICA), model parameters were fit using the training data for all neurons. Then, to determine how well each model was able to capture the activity of a single neuron using the testing data, we used the activity of all other neurons to calculate the activity of the latent variables (by setting the encoding weights of the left-out neuron to zero). We then performed a simple linear regression using the activity of the latent variables to predict the activity of the left out neuron. The *R*^2^ values reported are those obtained by comparing the true activity 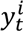 of neuron *i*at time *t* with the activity predicted by the various methods 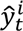 in this leave-one-out fashion, and averaging over neurons:

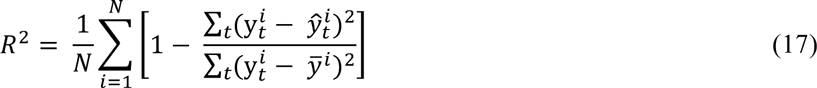

where 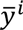 is the average activity of neuron *i*. Note that for this leave-one-out procedure, if just a single neuron is contributing to the activity of a latent variable, this procedure will result in a small *R*^2^ value for that neuron during cross-validation.

For simulated data, we can compare the true and inferred latent variables to ensure that an inferred latent variable isn’t driven by just a single neuron. Therefore, the *R*^2^ values reported for the simulated data are not calculated in this computationally-intensive leave-one-out manner, but rather use encoding and decoding matrices learned from the training data to compute the latent variables and the predicted activity. The resulting predicted activity is then compared to the true activity using eq. (17).

### Simulated data generation

We evaluated the performance of the RLVM using simulated datasets, which were generated using five non-negative latent variables that gave rise to the observed activity of 100 neurons. Note that these choices reflect our core hypotheses of the properties of latent variables in the cortex, and also match the assumptions underlying the RLVM model structure. Latent variables were generated by creating vectors of random Gaussian noise at 100 ms time resolution and then smoothing these signals with a Gaussian kernel. To enforce the non-negativity constraint on the latent variables, a positive threshold value was subtracted from each latent variable, and all resulting negative values were set to zero. Correlations between different latent variables were established by multiplying the original random vectors (before smoothing) by a mixing matrix that defined the correlation structure. Although smoothing and thresholding the correlated latent variables changed the correlation structure originally induced by the mixing matrix, the new correlation structures obtained by this procedure were qualitatively similar to those seen in experimental data.

The latent variables thus obtained acted as inputs to a population of neurons (Fig. 1). To calculate the coupling weights between the latent variables and the neurons in the population, a coupling matrix was created to qualitatively match the coupling matrices found in experimental data (compare Figs. 3B and 1C). Since the experimental data used later in the paper comes from a 2-photon imaging experiment, we chose to simulate data resembling 2-photon fluorescence traces. To compute simulated fluorescence traces for each neuron, first the firing rate of the neuron was computed as the weighted sum of the latent variables, with the weights defined in the coupling matrix. The resulting firing rate was used to produce a spike train using a Poisson spike generator. The spike train was then convolved with a kernel to create the calcium signal and finally Gaussian random noise was added to generate a simulated fluorescence signal.

**Figure 1.**
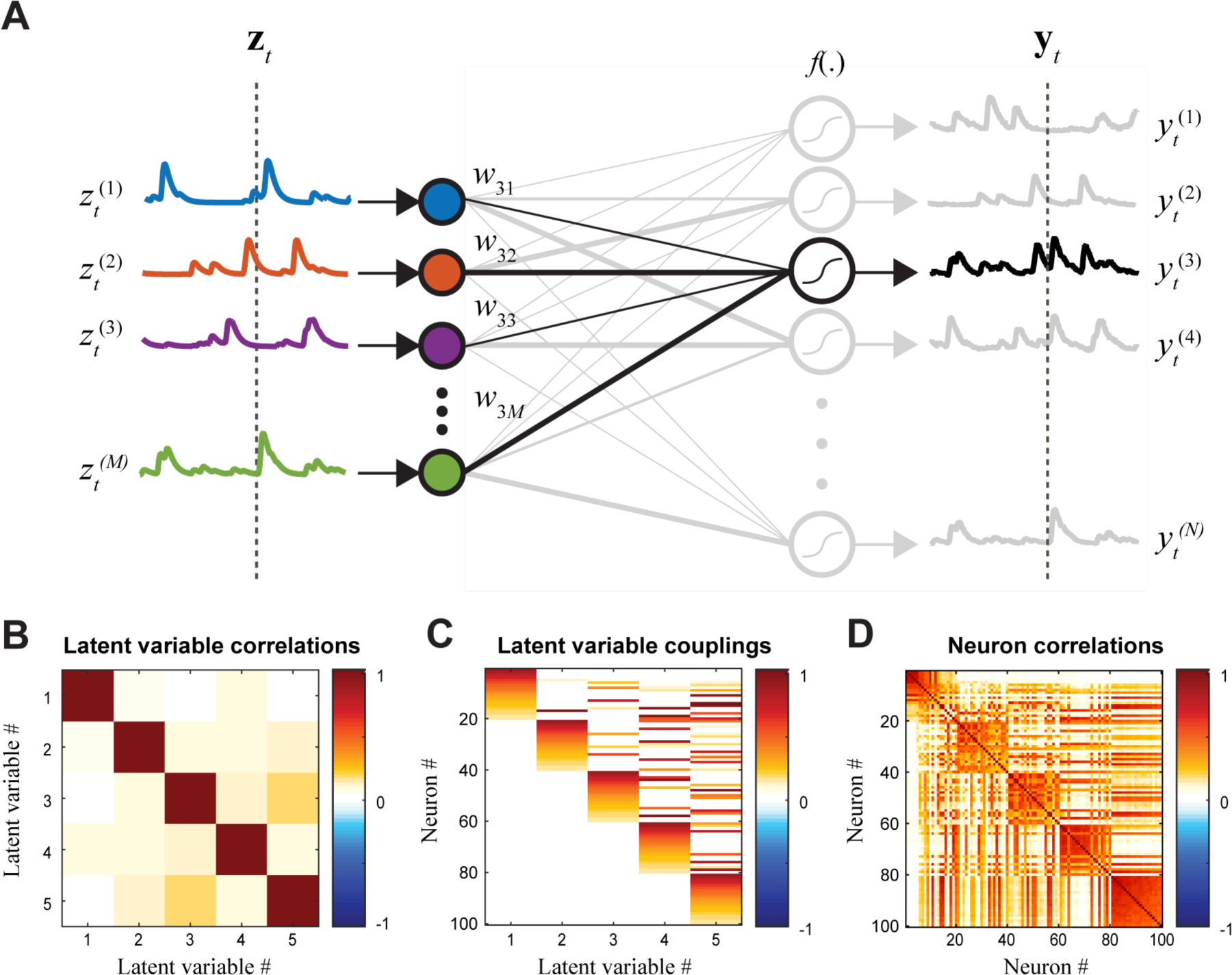
RLVM structure. The RLVM predicts the observed population response 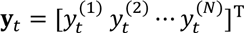 at a given time point *t* (dashed lines, *right*) using a smaller number of non-negative latent variables 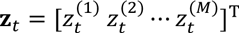 (dashed lines, *left*). The latent variables are weighted by a matrix W such that *w*_*ij*_ is the weight between neuron *i* and latent variable *j*, and the resulting weighted inputs are summed and passed through a nonlinearity *f*(.). There are additional offset terms for each neuron, not pictured here. (**B-D**) The hypothesized structure of the cortical network motivating the RLVM formulation is used to generate synthetic data, using five latent variables. **B.**Factors underlying cortical activity will often be correlated with each other, and our simulation of cortical activity used the correlation matrix shown between latent variables in generating simulated activity. **C.**The weight matrix between latent variables and each neuron, generated to approximate the coupling matrices found with experimental data (compare to Fig. 3B). **D.** The measured pairwise correlation matrix between each neuron, computed from simulated data. The correlations predicted by the RLVM arise solely from shared latent variable input and their correlations with each other, rather than pairwise coupling.

### Experimental data

#### Experimental protocol and data preprocessing

We evaluated the RLVM on data from the Svoboda lab (Peron et al., 2015), which has been made publicly available at http://dx.doi.org/10.6080/K0TB14TN. In this experiment mice performed a tactile discrimination task with a single whisker. During a given trial, neural activity was recorded from layers 2/3 of barrel cortex using 2-photon imaging with three imaging planes set 15 µm apart. These imaging planes constituted a subvolume, and eight subvolumes were imaged during a given session. Furthermore, those same subvolumes were imaged across multiple experimental sessions, and the resulting images were later registered so that activity of individual regions of interest (ROIs) could be tracked across the multiple sessions. Raw fluorescence traces were extracted from each ROI and neuropil corrected. For each ROI a baseline fluorescence *F*_0_ was determined using a 3-minute sliding window and used to compute Δ*F*/*F* = (*F*-*F*_0_)/*F*_0_.

#### Data selection

The publicly available dataset contains the Δ*F*/*F* fluorescence traces of tens of thousands of neurons imaged over multiple sessions for eight different mice. In order to select subsets of this data for analysis with the latent variable models, we restricted our search to volume imaging experiments where somatic activity was imaged in trained mice. We then looked for subsets of simultaneously imaged neurons that maximized the number of neurons times the number of trials imaged, selecting nine different sets of imaged neurons, three sets from each of three different mice. Within each set, neurons were removed from this selection if they met one or both of the following criteria: (1) more than 50% of the fluorescence values were missing (indicated by NaNs); (2) the fluorescence trace had a signal-to-noise ratio (SNR) less than one. To estimate the SNR, a smoothed version of the fluorescence trace was estimated with a Savitzky-Golay filter (using MATLAB’s built-in function `sgolayfilt`). The noise was estimated using the residual between the original trace and the smoothed trace. The SNR was then calculated as the variance of the smoothed trace divided by the variance of the noise. Furthermore, we removed trials from the data selection if NaN values existed in the whisker measurements or in one or more of the remaining fluorescence traces. See Table A1 in the Appendix for more information about the specific subpopulations of neurons analyzed.

#### Alignment of fluorescence traces across sessions

As described above, data from each experiment we used consisted of imaging the population activity over several recording sessions. Although fluorescence traces for each neuron were corrected for different baseline fluorescence levels in the online dataset, we found it necessary to recalculate session-specific baseline fluorescence levels in order to concatenate traces across different sessions. [Unlike the analyses in the original work (Peron et al., 2015), the models considered here were particularly sensitive to this baseline level because all fluorescence traces were analyzed jointly.] In the original work, baseline fluorescence level was calculated using the skew of the distribution of raw fluorescence values, under the assumption that more active neurons will have more highly skewed distributions. However, this monotonic relationship breaks down for very active neurons, whose distributions are not as skewed since there are very few values near the baseline level. Because we found many neurons in the dataset that fell into this last category, we recalculated baseline fluorescence levels on a session-by-session basis.

Using basic simulations of how the distribution of fluorescence values of a Poisson neuron depends upon its mean firing rate and SNR, we could match this with the data from each neuron to unambiguously infer its baseline fluorescence level. Specifically, for each neuron and each session, we measured both the SNR of its fluorescence (described above) and also measured the skewness of its distribution of fluorescence (using MATLAB’s built-in function `skewness`). We simulated neural activity with the same SNR while varying the mean firing rate until the resulting distribution of values matched the measured skewness. Once the optimal mean firing rate was determined, we could then use the simulation to determine the best estimate of the baseline fluorescence level on a session-by-session basis. This procedure led to improved model estimation for all latent variable methods.

#### Sorting of coupling matrices

The ordering of simultaneously recorded neurons is arbitrary, so we chose the ordering for the display of the coupling matrices W to highlight groups of neurons that share similar coupling patterns. We first sorted the rows using the coupling weights to the first latent variable (first column) for all neurons with a weight higher than a predefined threshold value. We then sorted all remaining neurons with coupling weights to the second latent variable (second column) above the same threshold. This procedure is repeated for each column of W, and produces a block diagonal structure (e.g., Fig. 3B). The last column is sorted without using the threshold so that it contains all remaining neurons.

## RESULTS

### Model formulation

The goal of latent variable modeling is to describe the activity of many simultaneously recorded neurons using a small number of latent variables. Consider a population of *N* neurons, with the ensemble of observed activity at time *t* represented by a vector **y**_*t*_ – this observed activity could, for example, be spike counts from multielectrode recordings or fluorescence values from 2-photon microscopy. The *M* latent variables will also have a vector of **z**_*t*_ at each time point, where *M* is a free parameter of the model (Fig. 1). The RLVM then attempts to predict the population activity **y**_*t*_ as a function *f*(.) of a linear combination of the latent variables **z**_*t*_

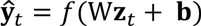

where W is a matrix of weights that describes how each neuron is coupled to each latent variable, and b is a vector of bias terms that account for baseline activity. For 2-photon data, it is appropriate to use a linear function for *f*(.), while for spiking data one can use a function that results in non-negative predicted values to match the non-negative spike count values (Paninski, 2004).

The vector of latent variables **z**_*t*_ will in principle represent all factors that drive neural activity, including both stimulus- and non-stimulus-locked signals. These factors may or may not be related to observable quantities in the experiment. For example, they could be related to “external” observables like motor output (Churchland et al., 2012) and pupil dilation (Vinck et al., 2015), or “internal” observables like the LFP (Cui et al., 2016), population rate (Schölvinck et al., 2015) or dopamine release (Schultz et al., 2015). However, while latent variables might be related to experimental observables, here we make no assumptions on such relationships in determining them.

The non-negative assumption on these latent variables is a key distinction between the RLVM and other latent variable models. It is motivated by the observation that any neural process, whether excitatory or inhibitory, must be realized using the action potentials or firing rates of neurons, neither of which can take on negative values. As a result, each latent variable is either inactive (zero) or active to varying degrees, which furthermore allows us to interpret the influence of a latent variable on a neuron as either excitatory (positive coupling weight) or suppressive (negative coupling weight). Additionally, such an assumption on the model structure breaks the rotational degeneracy characteristic of most linear models. This degeneracy arises in these models because the two solutions W**z**_*t*_ and (WU^T^)(U**z**_*t*_) are equivalent for any orthogonal matrix U, since U^T^U = I, the identity matrix. The RLVM will thus be constrained to find the best solution composed of non-negative latent variables. Notably, this assumption would hurt the ability of the model to describe population activity generated by unconstrained variables; however, if the population activity is truly generated by non-negative variables, this assumption should not impact model performance, and will likely contribute to an increased resemblance of the latent variables identified by the RLVM to underlying unobserved factors contributing to cortical activity.

A second key innovation of the RLVM is how it is fit to datasets. Inferring the time course of latent variables is challenging because of their high dimensionality, which has a different value for each time point in the experiment. Fitting the model using random initializations for both the latent variables **z**_*t*_ and model parameters {W, **b**} does not achieve good results because the optimization becomes stuck in poor local minima given such a high dimensional space. As a result, we used an autoencoder framework (Bengio et al., 2013) to fit all model components simultaneously, which provided reasonable initializations for the full fitting algorithm. The autoencoder optimizes both **z**_*t*_ and {W, **b**} by minimizing the mean square error (or any appropriate cost function) between the true activity and the activity predicted by eq. (13). By using these values to initialize the MML algorithm we were able to achieve much more accurate solutions, detailed below.

### Validation of the RLVM using simulated data

In order to understand the fitting behavior of the RLVM and ensure that it performs as expected, as well as to compare the RLVM with other latent variable models, we generated simulated data with five latent variables and 100 neurons. This data was generated under the assumption that the input to each neuron comes from a weighted combination of a small number of correlated, non-negative latent variables. The resulting input to each neuron was then passed through a spiking nonlinearity to produce its firing rate, which was then used to randomly generate spike counts using a Poisson process. Since the experimental data used later in the paper comes from a 2-photon imaging experiment, simulated data was further processed by convolving the generated spike trains with a kernel to simulate calcium dynamics, and finally adding Gaussian noise.

#### Evaluation of RLVM fitting methods

We first demonstrate the importance of the autoencoder stage of our optimization procedure, the results of which are used as to initialize the MML algorithm, which is then used to optimize the model parameters and latent variables (see Methods). To quantify the goodness of fit for each model type (random initialization vs. autoencoder initialization) we calculated the Pearson correlation coefficients (*r*) between the true and inferred latent variables. Using random initializations for this procedure led to poor solutions for the latent variables (*r* = 0.781 ± 0.020; mean *r* ± SEM over 20 initializations), whereas first fitting the autoencoder (itself initialized randomly) and using the resulting parameters to initialize the MML algorithm led to far more accurate solutions (*r* = 0.971 ± 0.001). The superior results achieved by initializing with the autoencoder solution are due to the high dimensionality of the problem – in the MML algorithm employed here there are relatively few constraints imposed on the latent variables (non-negativity and some degree of smoothness, see Methods), which results in many local minima. In contrast, the latent variables of the autoencoder are constrained to be a linear combination of the recorded population activity, and this constraint results in a much smaller space of model solutions.

In fact, we found that the latent variables resulting from the autoencoder-initialized MML optimization were extremely similar to the initial values found by the autoencoder itself (*r* = 0.994 ± 0.000). The main difference between these solutions is that the additional optimization step following the autoencoder smooths the time course of the latent variables, whereas the autoencoder latent variables are not generally smooth. Due to the fact that the latent variables from the autoencoder do not have to be separately inferred for cross-validation data, it is often convenient to use the autoencoder solutions themselves, forgoing the MML optimization altogether. This has the added advantage here in making model performance more straightforward to compare against the other latent variable models.

Because there are many factors that can affect the performance of the autoencoder network used to fit the RLVM, we also looked at the effects of varying parameters governing both the simulated data generation and the fitting procedure to better understand the autoencoder’s sensitivity to these variables. We found that the autoencoder can accurately recover the latent variables and coupling matrix even with small amounts of data (Fig. A1A) and low SNR (Fig. A1B). We explored the sensitivity of the autoencoder to different values of the regularization parameter on the encoding and decoding weights (λ_1_ and λ_2_, respectively, in eq. 16), and found that the results obtained by the autoencoder are constant across several orders of magnitude (Fig. A1C). In practice, we also found that the autoencoder solutions were not prone to getting stuck in different local minima given random initializations of the autoencoder parameters. These experiments suggest that the autoencoder is a robust fitting method for the RLVM that does not need large amounts of data or precise tuning of optimization parameters in order to produce accurate results.

We also tested whether the RLVM’s non-negativity constraint is essential for recovering the correct latent variables from the simulated data. Again using *r* as a goodness-of-fit measure for the inferred latent variables, we fit the RLVM to the simulated data (using the autoencoder) with different functions for g(.) in eq. (12). We found that using the rectified nonlinearity (ReLU function) led to much more accurate solutions (*r* = 0.963 ± 0.002; ± mean *r* ± SEM over 20 initializations) than when using a non-rectified (linear) version of the RLVM (*r* = 0.573 ± 0.021). This illustrates the importance of using the nonlinearity to enforce the non-negativity of latent variables, in order for the RLVM to recover the latent variables generated with such a non-negative constraint.

#### Comparison of RLVM with other latent variable methods

To understand how the RLVM compares with other latent variable methods, we also fit PCA, FA and ICA models to the simulated data (Fig. 2). Such simulations are useful for putting in context results from real data, where the ground truth is not known. We first compared the latent variables inferred by the different models (Fig. 2A), again using *r* as a goodness-of-fit measure. The RLVM and FA perform extremely well, while PCA and ICA perform rather poorly (Fig. 2B). The good performance of the RLVM should be unsurprising, given that the data was generated according to the assumptions of the RLVM. The fact that FA performs so well is perhaps more surprising given that it assumes independent Gaussian variables, but these assumptions are only used for determining the initial coupling matrix; the final coupling matrix is determined using varimax rotation (MATLAB default), and the resulting latent variables are determined using linear regression (MATLAB default), which makes no assumptions about their distribution. PCA and ICA do not infer the correct latent variables because they make assumptions about the latent variables being uncorrelated (PCA) or independent (ICA), neither of which is true of the simulated data (Fig. 2B).

**Figure 2.**
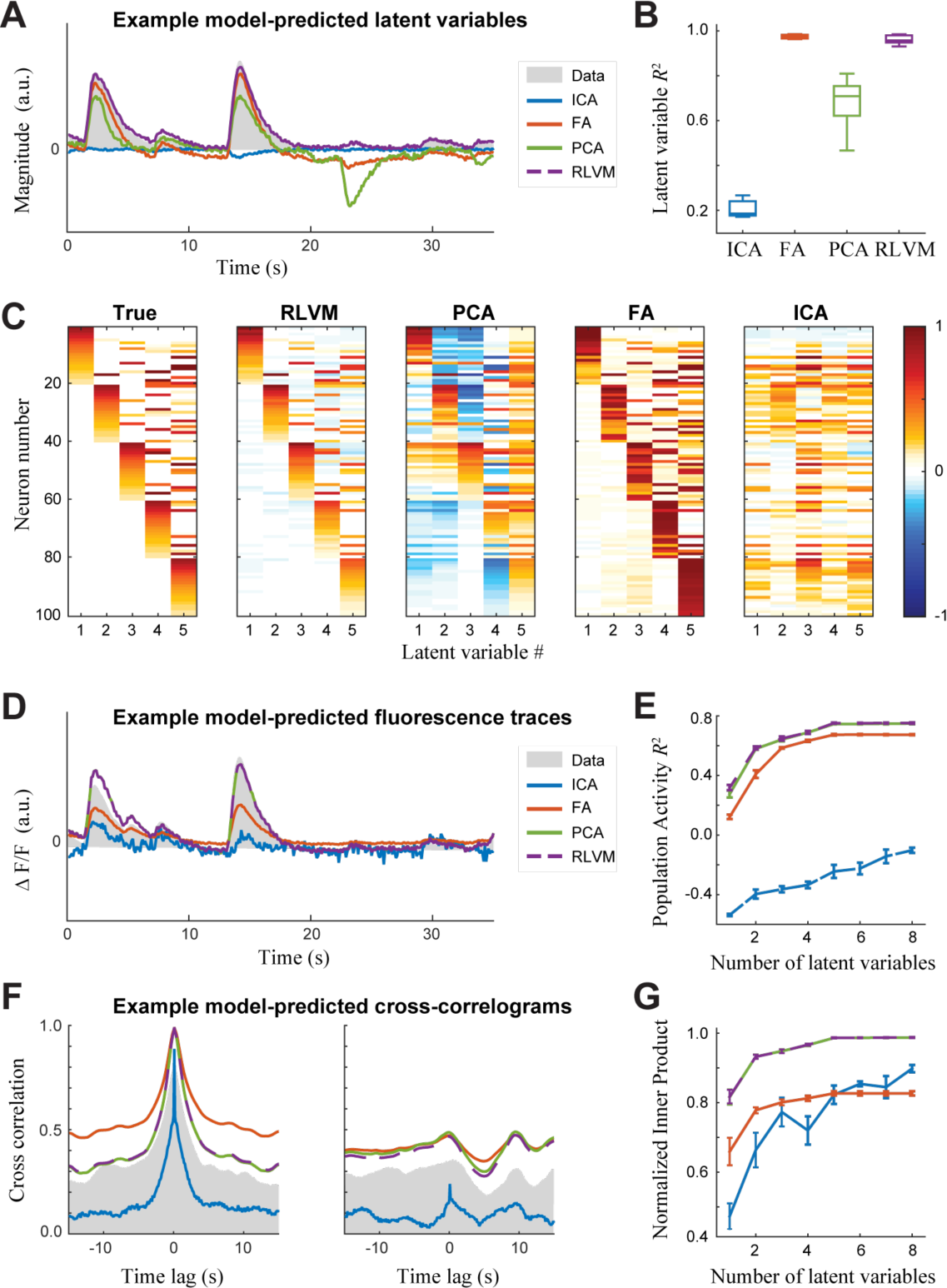
Comparisons between latent variable methods applied to simulated data. Four different latent variable methods were fit to simulated data (see Fig. 1B-D), and cross-validated model performance measures were computed. **A.** The time course of a representative latent variable, compared to predictions inferred from each method. Note that the FA and RLVM methods are both highly overlapping with the true latent variables. **B.** Correlation coefficient between true latent variables and those inferred by each method, demonstrating the superior performance of the FA and RLVM methods. Boxplot shows the distribution of correlation coefficients over latent variables and cross-validation folds. **C.** The matrices of coupling weights between neurons and latent variables, inferred by each method. For comparison, the coupling matrix used to generate the simulation is shown (*left*, reproducing Fig. 1C). **D.** Representative fluorescence trace of one neuron from the simulated data compared to the predicted trace from each method. Despite their performance in predicting the latent variables (A), here FA does poorly, and PCA does well, as does the RLVM. **E.** Median *R*^2^ value across neurons between true fluorescence traces and those predicted by each method over the simulated neurons, plotted versus the number of latent variables specified during the fitting procedure. Error bars indicate the standard deviation of the mean of median *R*^2^ values over cross-validation folds. Model performance in each case plateaus for the true number of latent variables, but is limited in each method due to how the neural activity is generated. **F.** Two example cross-correlograms from simulated neuron pairs plotted with the corresponding cross-correlograms calculated from predicted traces for each method. **G.** The ability of each method to reproduce the pairwise cross-correlations between neurons, measured as the normalized inner product between the true correlation matrix and those calculated from predicted traces for each method. This is again plotted against number of latent variables specified for each method, and plateaus at the true number of latent variables. Error bars indicate standard deviation of the mean inner product over cross-validation folds.

Given the inferred latent variables, we were also interested in how well each method captured the coupling weights between these latent variables and each neuron. Visual inspection of the coupling matrices (Fig. 2C) shows that the RLVM and FA performed much better than PCA or ICA, due to the accuracy of their inferred latent variables. Again, the strong assumptions that PCA and ICA place on the latent variables prohibits their accurate estimation of the coupling matrices.

For all four models considered here, the predicted activity of an individual neuron is given by the sum of the latent variables (Fig. 2A, B), weighted by the values in the proper row of the coupling matrix (Fig. 2C). To quantify this prediction accuracy, we used the coefficient of determination (*R*^2^; see Methods) between the true and predicted activity (Fig. 2D, E). Interestingly, even though the RLVM and FA produced similar latent variables and coupling matrices, FA did not predict the population activity as well as the RLVM. This was mostly due to many large weights in the FA coupling matrix (compare the red diagonal blocks in Fig. 2C to the true coupling matrix), which is an artifact of the varimax rotation step common in many FA algorithms.

Perhaps surprisingly, PCA performed just as well as the RLVM, even though PCA does not infer the correct latent variables or estimate the correct coupling weights. The reason for this is that the RLVM and PCA both minimize the reconstruction error in their cost function (explicitly and implicitly, respectively); however, because PCA does not constrain the latent variables to be positive, it reconstructs the population activity using both positive and negative values. This leads to differences in the latent variables (Fig. 2A) and coupling matrices (Fig. 2C), but can result in an equivalent prediction of activity (Fig. 2D). This difference between the RLVM and PCA in their descriptions of the population activity is a crucial point that we will return to when evaluating the PCA solutions on real data.

Finally, we evaluate each method on its ability to account for observed correlations between neurons. Many previous approaches have focused on explaining pairwise correlations directly (Schneidman et al., 2006; Pillow et al., 2008; Ohiorhenuan et al., 2010), which requires parameters for each pair of neurons. However, as we see from our simulation, even just five latent variables can produce a complex pattern of pairwise interactions (Fig. 2F). Thus, latent variable methods offer the ability to explain such correlations using many fewer parameters. To quantify each model’s ability to capture these correlations, we compare the cross-correlogram at the zero-time-lag point between data and prediction from each neuron pair (which forms the correlation matrix). This agreement was measured using the overlap between the true correlation matrix and the predicted correlation matrix (Fig. 2G). The results mostly mirror the ability of each method to predict the population activity (Fig. 2E), with the RLVM and PCA capturing more of the correlation structure than FA and ICA.

### Application of the RLVM to 2-photon experiments in primary somatosensory cortex

With the RLVM validated for simulated data, we next apply the RLVM to the experimental dataset from Peron et al. (2015). We selected this dataset because it involves a complex task with many “observables” related to behavior and task context, many of which are outside of direct experimental control, but likely related to cortical activity. Additionally, there were a large number of neurons recorded over long periods of time, which allowed us to test the RLVM on a dataset of appreciable size. In this experiment, mice performed a pole-localization task, in which a pole was lowered at a distal or proximal location next to the animal’s snout. The animal could touch the pole with a single whisker, and then had to signal its decision after a delay period by licking one of two lick-ports following the onset of a brief auditory cue. For the analyses in this work, we used a particular subset of this data (see Appendix, Table A1) selected based on the size of the neural population imaged, the length of time imaged, and its signal-to-noise ratio (see Methods).

We first determined how well the different latent variable models predicted the observed population activity (Fig. 3A) (see Methods for model-fitting procedures). The relative performance of the methods is similar to their performance on the simulated data (Fig. 2E). For the RLVM, PCA and FA, there was at first a rapid increase in prediction performance as the number of latent variables increases, and then the performance began to plateau between five and ten latent variables. While this plot does not directly indicate how many “true” latent variables generate the data, it is important to note that relatively few are needed before the performance plateaus. Because there is no explicit point where this occurs, we selected a point where there was only a marginal increase in performance (six latent variables) for all subsequent analyses.

**Figure 3.**
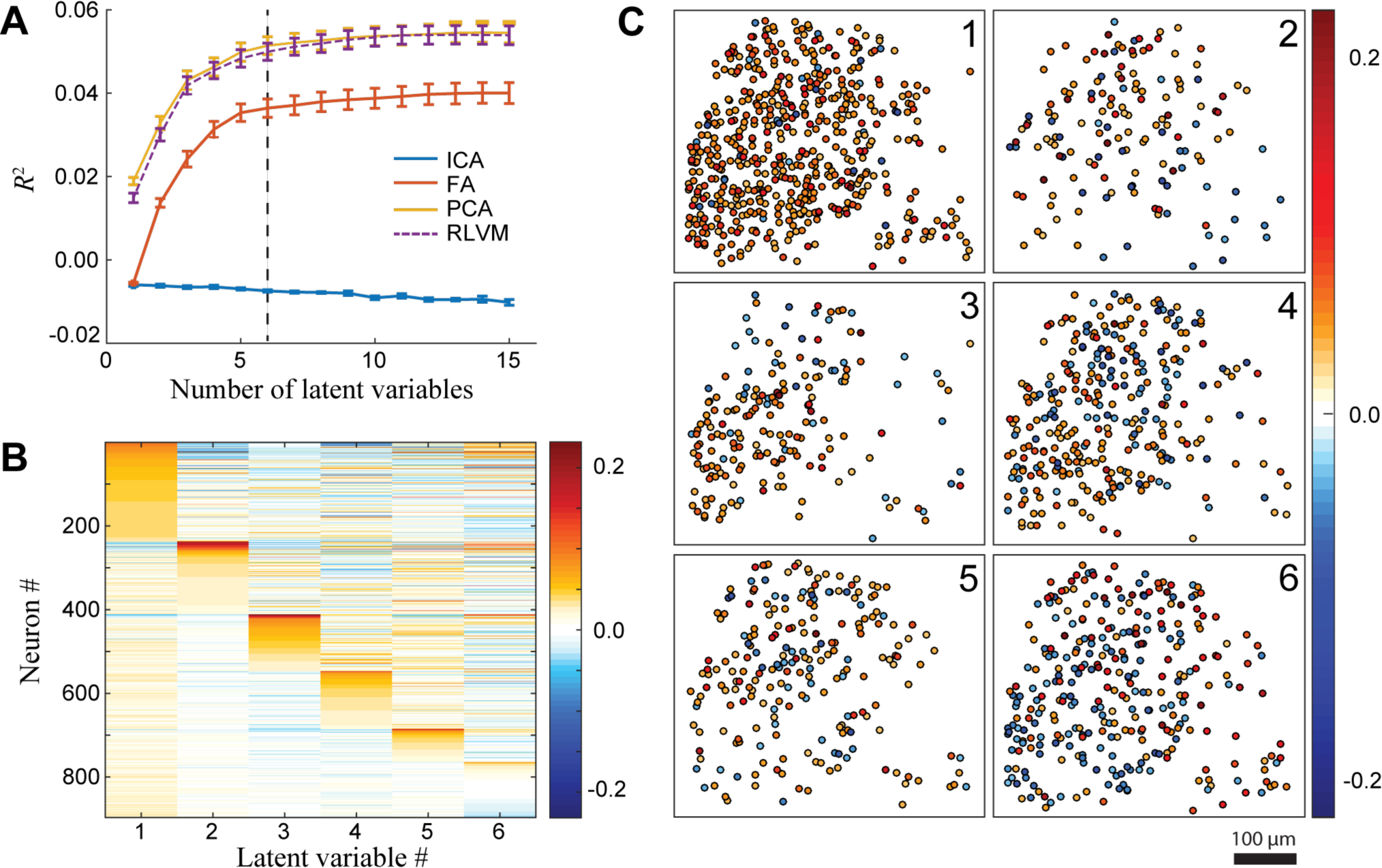
Latent variable methods applied to two-photon imaging of mouse somatosensory cortex. **A.** The ability of the models to reproduce the observed data depends on the number of latent variables used, measured by *R*^2^ between the measured and predicted activity. The relative performance of the different methods is ordered like their applications to the simulated data (Fig. 2D). Because there was no clear saturation point of the *R*^2^ values, models with six latent variables (dashed line) were used for subsequent analyses. **B.** The coupling weights of the RLVM between each neuron and each latent variable, ordered to make visualization of neuron clusters in each latent variable easier (see Methods). **C.** The spatial patterning of coupling to each latent variable is pictured by displaying neurons whose coupling strength is greater than 15% of the maximum coupling strength for the latent variable, and color-coded to show the magnitude of this coupling. The imaged neurons were within a single barrel (of mouse primary somatosensory cortex), and the coupling to latent variables exhibited no clear spatial pattern.

Once the latent variables are determined, the coupling matrix of the RLVM allows us to understand how each neuron combines these variables to produce its predicted activity (Fig. 3B). One of the advantages of using 2-photon data is that it provides the spatial locations of the neurons, and we can use that information to determine if there is any spatial structure in the coupling weights to the latent variables. We plotted a subset of the weights (any with an absolute magnitude greater than 15% of the maximum absolute magnitude for each latent variable) according to each neuron’s spatial location (Fig. 3C). The positive and negative weights are intermingled in these plots, and no discernible spatial structure exists. This is expected in part because these neurons are imaged within a single barrel, and thus all belong to a single cortical column. Nevertheless, this illustrates how latent variables can provide a new way to investigate the functional organization of cortex.

With simulated data we were able to directly compare the latent variables inferred by each method with the ground truth, but with experimental data we have no direct way to validate the latent variables that each method detected. Instead, we hypothesized that latent variables will be related to factors that might be directly observed in the experiment. We thus begin by comparing the time course of latent variables discovered by the RLVM to different elements of the experiment. In this case, there were four “trial variables” measured in this dataset: the timing of whisker touches against the pole, the onset of the auditory cue that signals the animal to make its choice, the onset of reward delivery when the animal makes the correct choice, and the timing of licks. Clearly, the time courses of different latent variables had some relationship with some of the trial variables (Fig. 4A).

**Figure 4.**
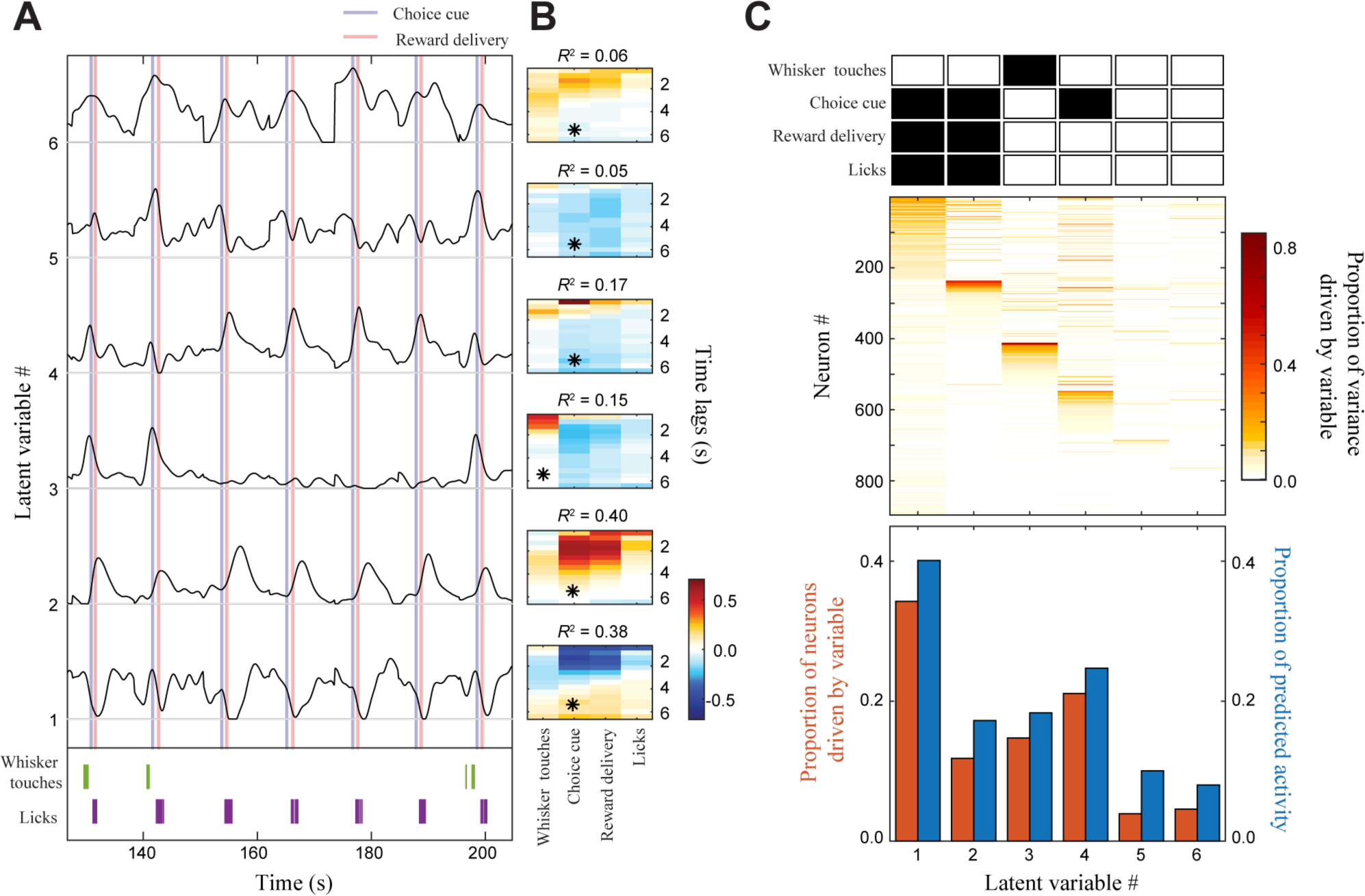
Relationship of the latent variables inferred by the RLVM to experimentally observed trial variables. **A.** An 80-second sample of the predicted activity of six latent variables (extending over eight task repetitions), demonstrating the relationship of some of these latent variables to trial variables observed during the experiment: the auditory cue that signals the animal to make its choice (blue vertical), the onset of reward delivery when the animal makes the correct choice (red vertical), the timing of whisker touches against the pole (*bottom*, green), and the timing of licks (*bottom*, purple). Latent variables are ordered (*bottom* to *top*) based on the magnitude of their variance. **B.** Quantification of the relationship between the latent variables and trial variables, where the values of each trial variable at different time lags were used to predict the activity of each latent variable using linear regression. The ability of the trial variables to predict the latent variable time course (measured by *R*^2^) was used to assess the relationship between the two. The trial variable with the largest *R*^2^ value is marked with an asterisk, and the corresponding *R*^2^ value is displayed above the coefficients. **C**. *Top*: The shaded boxes indicate which trial variables are capable of predicting each latent variable. *Middle*: The fraction of measured activity of each neuron accounted for by each latent variable. The resulting matrix is related to a weighted version of the coupling matrix (Fig. 3B), and demonstrates the relative contribution of each latent variable to the observed population activity. *Bottom*: The fraction of observed neurons driven by each latent variable (red), and the relative fraction of predicted neural activity explained by each latent variable (blue).

To quantify these observations, we used linear regression in order to predict the activity of each latent variable using the four observed trial variables. We performed a separate linear regression for each trial variable, which did not take into account the correlations that exist among the trial variables, like reward delivery and licks. This leaves open the possibility that a latent variable is actually driven by reward delivery, but is equally well-predicted by licks because of the tight correlation between these two variables. There is also the possibility, however, that the animal licks many times when the reward is not delivered (such as on error trials) and so we considered all trial variables independently. Furthermore, coefficients for the linear regression include lagged time points, which allowed the regression model to capture the extended temporal response of fluorescence transients (Fig. 4B). We found that latent variables #1, #2 and #4 are well-predicted by the reward portion of the trial, latent variable #3 is well-predicted by whisker touches, and latent variables #5 and #6, which don’t have any discernible trial-locked patterns, are not well-predicted by any of these four trial variables.

With these quantitative measures, we can label each latent variable with the set of trial variables that best predict it. To do so, we required that (1) the *R*^2^ value using that trial variable was greater than 0.10, and (2) the *R*^2^ value was greater than one-half the largest *R*^2^ value among all trial variables. If both these conditions were met, we considered the latent variable to be driven (though perhaps not exclusively) by that trial variable (Fig. 4C, lower panel). We found that, even though this population of neurons is located in the primary somatosensory cortex, only one of the latent variables is identified with whisker touches, while three of the latent variables are identified with the reward portion of the trial.

Besides just knowing which trial variables are correlated with each latent variable, we were interested in quantifying how strongly each latent variable influences the population response. First, we looked at how each latent variable contributes to the overall proportion of predicted activity. To do this for a given latent variable, for each neuron we calculated the variance of the latent variable, weighted by the neuron’s coupling strength to that latent variable (from the matrix in Fig. 3B). This value was divided by the variance of the neuron’s total predicted activity, and the resulting value is a measure of how much that latent variable contributes to that neuron’s predicted activity. [Note that, because latent variables can be correlated, these proportions will not add to one since we ignored cross-covariances.] These values were then averaged over all neurons to obtain a measure of the proportion of predicted activity driven by the given latent variable (Fig. 4C, upper panel, blue bars). Latent variable #1, which is driven by the reward portion of the trial, contains the largest proportion of predicted population activity. Latent variable #3, which is the only latent variable identified with the stimulus, contains the third largest proportion, while latent variables #5 and #6, which are not identified with any trial variables, contain the lowest proportion of predicted activity.

We also quantified how strongly a latent variable influences the population by measuring the proportion of neurons driven by that latent variable. To quantify this for a given latent variable, for each neuron we divided the variance of the weighted activity of the latent variable by the variance of the neuron’s measured activity, smoothed using a Savitzky-Golay filter to remove noise variance (implemented with the MATLAB function `sgolayfilt`) (Fig. 4D). We then considered a neuron to be driven by that latent variable if the proportion of total measured activity exceeds 0.10 (Fig. 4C, upper panel, red bars). Similar to previous observations, latent variable #1 affects the largest proportion of neurons, at about one-third, while latent variables #5 and #6 affect the smallest proportions of neurons.

We performed the same analyses as in Fig. 4 using PCA to see if there were fundamental differences in its description of the population activity (Fig. 5). The latent variables inferred by PCA (Fig. 5A) do in fact contain features that are correlated with the trial variables, but these features tended to be more mixed than in the RLVM latent variables. To illustrate this, consider RLVM latent variables #1 and #2, which are associated with suppressive and excitatory activity during the reward phase, respectively (Fig. 4A). While the RLVM cleanly separates these two subpopulations, they are mixed together in the first principal component of PCA (Fig. 5B, *middle*; neurons ~0-50 and ~240-260, respectively). PCA mixes these two subpopulations because such a combination into a single principle component explains the greatest amount of variance in the data, and this combination is possible because PCA is not restricted to using non-negative latent variables.

**Figure 5.**
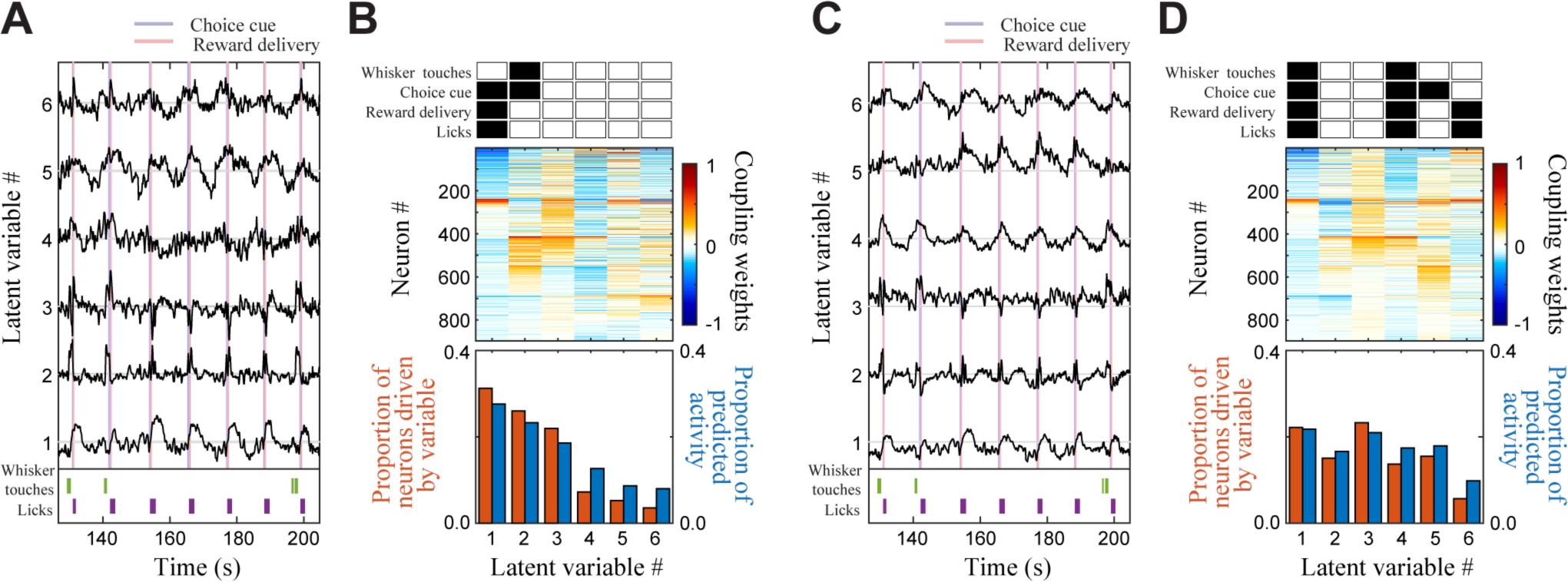
Latent variables inferred by PCA and a linear RLVM show a weaker relationship to individual experimental trial variables. PCA (A, B) and a linear RLVM (where latent variables were not constrained to be non-negative) (C, D) were fit to the same experimental data as in Fig. 4. **A.** Latent variable time courses inferred by PCA over the same interval as in Fig. 4A, ordered from *bottom* to *top* by variance explained. There is a clear mixing of information relative to the RLVM latent variable time courses (Fig. 4A). Latent variable #3, for example, has positive deflections aligned with whisker touches (similar to RLVM latent variable #3) combined with negative deflections aligned with the onset of the reward period (opposite sign relative to RLVM latent variable #4). **B.** *Top*: Shaded boxes indicate which trial variables are related to the latent variables. *Middle*: Coupling matrix between latent variables and each neuron (neurons are ordered the same as those in Fig. 4C). This illustrates how the first few principal components mix inputs from several sources, likely because PCA is based on explaining the greatest fraction of variance with each principal component rather than separating the underlying causes. *Bottom*: The summed influence of each latent variable on the population activity (matching measures in Fig. 4C, *bottom*). **C.** Latent variables inferred by a linear RLVM. **D.** Same measures as those calculated in B. Both the PCA and linear RLVM latent variables mix features from the RLVM latent variables, which is apparent in their coupling matrices (B, D, *middle*).

To determine if the non-negativity constraint on the RLVM is responsible for the differences between the PCA and RLVM solutions, we fit the RLVM on the same data without constraining the latent variables to be non-negative. The latent variables inferred by this non-rectified version of the RLVM were qualitatively similar to PCA’s latent variables (Fig. 5C), and indeed this model’s latent variables exhibit the same type of mixing as the PCA latent variables. This demonstrates that the RLVM’s ability to separate these subpopulations of neurons is mainly due to the rectified nonlinearity, rather than an artifact of PCA’s constraint that the latent variables must be uncorrelated. This result is similar to the earlier comparison between the RLVM and non-rectified version of the RLVM on simulated data, where we found that the rectified nonlinearity was crucial for inferring the correct latent variables.

This example also illustrates that although the RLVM and PCA are able to explain the same amount of population activity (Fig. 3A), the underlying latent variables can differ dramatically due to rectification (similar results were seen with FA – data not shown). This same result was seen in the simulated data (Fig. 2A), and suggests that – if population activity is indeed composed of non-negative latent variables – the structure of the RLVM makes it a more appropriate method for studying neural population activity.

To demonstrate that the above results from the RLVM (Figs. 3 and 4) are consistent across different populations of neurons and different animals, we repeated these analyses using three populations of neurons from each of three animals (see Table A1 in the Appendix for more detailed information). The nine populations contain anywhere from 106 to 896 neurons, and the prediction performance of the RLVM for each population is plotted in Fig. 6A. It is interesting to note that all of these curves mostly plateau before reaching 10 latent variables, despite the fact that the number of neurons in these populations spans almost an order of magnitude. To repeat the analyses in Fig. 4 we used six latent variables for each population (Fig. 6B). Values were calculated as before, but averaged over latent variables. (Averaging over all neurons in each latent variable, which takes into account the relative sizes of the populations, did not qualitatively change these results).

**Figure 6.**
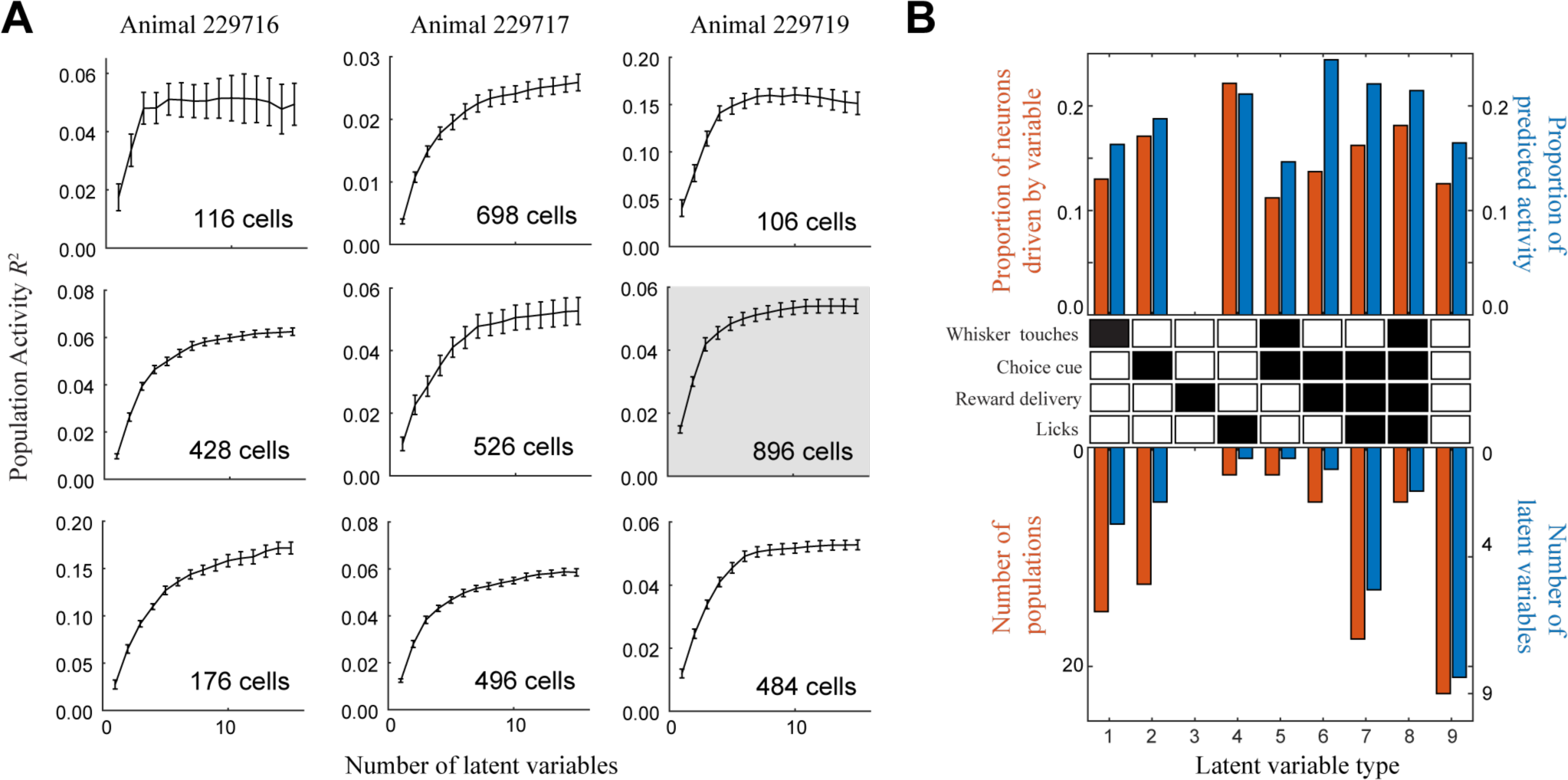
Consistent types of latent variables detected across experiments. **A.** *R*^2^ between the measured activity and the activity predicted by the RLVM for different imaged populations of neurons. The highlighted plot corresponds to the population of neurons analyzed in Figs. 3 and 4, and reproduces the RLVM values in Fig. 3A. Across experiments there is a similar dependence of *R*^2^ on the number of latent variables, although the overall magnitude of *R*^2^ values depends on the number of neurons and the level of noise in each experiment. **B.** *Top*: The amount of variability accounted for by each “type” of latent variable across all nine populations, using six latent variables per population (same measures as those calculated in Fig. 4C, averaged across the number of latent variables of each type). Even though all imaged populations were located in primary somatosensory cortex, across experiments much of the population activity was related to non-tactile sources. *Middle*: Latent variables are identified by the combination of trial variables each is related to (same criteria as those used in Fig. 4C). *Bottom*: Red bars indicate the total number of populations that contain at least one example of the latent variable type (out of nine total populations) and the blue bars indicate the total number of latent variables of each type (out of 54 total latent variables). Latent variable types identified with whisker touches (#1, #5 and #8) comprise only a small proportion of the latent variables, while latent variable types identified with the reward portion of the trial (#2-#4, #6 and #7) are much more prominent. Latent variables that were not identified with any trial variables (#9) were present in every population, and influenced the population activity to a similar degree as the others.

This meta-analysis shows that the results from Figs. 3 and 4 hold across different populations in different animals: the latent variables associated with the reward portion of the trial account for the largest proportion of the predicted activity in the populations; latent variables associated with the stimulus are found in the majority of populations, but drive a smaller proportion of neurons; and variables that are not identified with any trial variables are found in all populations, and tend to drive a smaller proportion of neurons. Together, these results (Figs. 3-6) demonstrate the usefulness of the RLVM as a tool for studying population responses in cortical areas, and suggest that latent variable models will be crucial to arriving at a deeper understanding of cortical function.

## DISCUSSION

Recordings of the activity from increasingly large numbers of cortical neurons provides opportunities to gain insight into the underlying factors driving cortical activity. Given that there are fewer variables underlying the activity than the number of neurons being recorded, latent variable approaches provide a way to infer the time course of these underlying factors and their relationship to neural activity. Here, we presented the Rectified Latent Variable Model (RLVM), which is unique in that it assumes a nonlinear structure on the network appropriate for neural activity – namely, that underlying factors are non-negative (rectified). The RLVM can be fit without needing to rely on a number of statistical assumptions characteristic of past latent variable models, such as the specification of particular distributions for the latent variables. The RLVM is robust to many aspects of data acquisition and model fitting (Fig. A1), and scales well with increasing numbers of neurons and recording length (Fig. A2).

The results of the simulated data experiments demonstrate that the RLVM is able to outperform PCA, FA and ICA across a variety of measures. It is able to properly recover the latent variables that generated the simulated data (Fig. 2B), as well as each neuron’s coupling weights to those latent variables (Fig. 2C). This guarantees that the method is able to predict single neuron activity well (Fig. 2E), which thus implies the method is able to accurately capture the structure of the pairwise correlation matrix (Fig. 2G).

Our results on experimental data demonstrate the utility of the RLVM as a tool for addressing questions about the structure of joint responses in large neural populations. Some of the latent variables inferred by the RLVM have clear relationships with measured trial variables, indicating that these latent variables have meaningful interpretations. We also demonstrated that the nonlinear nature of the RLVM leads to important distinctions in the description of the population activity when compared to a method like PCA, which has consequences for further understanding the role these latent variables play in cortical function.

### Relationships to other latent variable models

Latent variable models can be classified into two broad classes: static models, which do not take temporal dynamics of the latent variables into account, and dynamic models, which do. The RLVM has elements of both, although is more directly comparable to static models like PCA, FA and ICA. These models are known as linear factor models, so termed because there is a linear transformation from latent variables to predicted activity. While this need not be the case in the general RLVM framework (eq. 1), the formulation of the RLVM for 2-photon data uses this assumption as well. One advantage of the RLVM over these other linear factor models is that it does not specify any statistical constraints on the latent variables, which allows it to accurately capture correlated latent variables. Furthermore, due to the non-negativity constraint on the latent variables, the RLVM is able to identify latent variables that more closely resemble the form of expected inputs into the cortex, and does not have multiple equivalent solutions that arise from orthogonal transformations like some linear factor models.

In the absence of such nonlinearities, there is a close relationship between the RLVM and PCA. If the nonlinearities *f*(.) and *g*(.) in eq. (11) of the RLVM are linear, and the mean square error cost function is used, then the autoencoder solution of the RLVM lies in the same subspace as the PCA solution (Boulard and Kamp, 1989). The only difference is that the components of the RLVM can be correlated, whereas PCA requires them to be uncorrelated. However, using nonlinear functions for *f*(.) and/or *g*(.) allows the RLVM to capture more complex structure in the data than a linear model like PCA (Japkowicz et al., 2000).

The RLVM structure is also comparable to energy-based models, which is another class of static models that is exemplified by the Restricted Boltzmann Machine (RBM) (Koster et al., 2014). In the case that both the nonlinearities *f*(.) and *g*(.) in eq. (11) are sigmoids, the RLVM has the exact same mathematical structure as the standard RBM. Despite this similarity, the energy-based cost function of the RBM results in a model fitting approach that is significantly different from that of the RLVM. While standard RBMs are used with binary data, they can be extended to work with Gaussian distributions, in which case they closely resemble FA (Hinton, 2012).

The RLVM structure also contains elements of dynamic latent variable models, due to the log-prior term log *p*(**Z**) in eq. (2). The smoothing prior that we use here allows latent variable values at time points *t-1* and *t+1* to influence the value at time *t*. This is similar to the smoothing prior of Gaussian Process Factor Analysis (GPFA) (Yu et al., 2009), which allows a latent variable value at time point *t* to have a more complex dependence on past and future time points. However, as the name implies, GPFA is based on FA, and imposes similar statistical constraints on the latent variables that we try to avoid with the RLVM for reasons mentioned above. The state space models (Paninski et al., 2009) are another class of dynamic latent variable models, and constrain each latent variable at time *t*to be a linear combination of all latent variables at time *t-1*. Unlike the RLVM, which specifies a fixed relationship between the time points in the dynamics model, state space models fit the dynamics model to the data. This allows one to model the causal relationship between latent variables, but comes at the expense of making a strong assumption about the form of that relationship (namely, that latent variables are only determined by their values at the previous time step).

We found that the solutions for both the static and dynamic versions of the RLVM were similar, due in part to the simple dynamics model we imposed. However, the nature of 2-photon data does not lend itself to more restrictive dynamics models (like the state space models) because of the slow time scales. The investigation of more complex dynamics models in the RLVM is a direction for future work.

### Model extensions

The RLVM is a flexible model framework because there are many possible extensions that can be incorporated, depending on the desired application area. For example, additional regularization penalties can enforce desired properties on either the coupling weights or the latent variables. An L1-norm penalty on the coupling weights can enforce sparseness, so that each neuron only receives nonzero input from a small number of latent variables. An L_1_-norm penalty can likewise be applied to the activities of the latent variables, so that the activity of each latent variable is sparse.

One limitation of the RLVM is that it is only able to model additive interactions between the latent variables. Although there is evidence to support the existence of additive interactions in cortex (Arieli et al., 1996), and although they are commonly used for modeling (Okun et al., 2015; Schölvinck et al., 2015; Cui et al., 2016), there has been recent interest in modeling multiplicative interactions (Lin et al., 2015; Rabinowitz et al., 2015). It is theoretically possible to extend the RLVM to model non-additive interactions by adding more hidden layers to the model. This approach effectively allows a neural network to transform the latent variables into the observed data in a nonlinear manner, and is the basis of “stacked” autoencoders (Bengio et al., 2013), which we leave as a direction for future work.

For some analyses, it may not be desirable to have the effects of the stimulus represented in the activity of the latent variables. In this case it is possible to incorporate a stimulus model into the RLVM, such that the activity of each neuron is driven by the weighted sum of the latent variables plus terms that capture the stimulus response. These stimulus terms could be a simple, nonparametric PSTH-based model of the stimulus, or involve a more complicated parametric form (McFarland et al., 2013). Regardless of how the effects of the stimulus are captured, using the autoencoder variant the RLVM can still be fit using a standard gradient descent algorithm, and allows for the investigation of the relationship between stimulus processing and ongoing cortical activity.

The recent development of new recording technologies like high-density multi-electrode arrays and 2-photon microscopy is leading to increasingly large and rich neural datasets. We’ve shown here that the RLVM can be used effectively to analyze 2-photon datasets, and it is also possible to apply this model to spiking data by using a negative log-likelihood cost function that assumes Poisson noise (Fig. A4). The RLVM is thus a simple and extendable model that can be used to analyze both types of large population recordings, and in doing so can help uncover neural mechanisms that may not be obvious when studying the responses of single neurons.

## ACKNOWLEDGEMENTS

The authors thank the Svoboda lab and the CRCNS database for providing the mouse barrel cortex data. We also thank J. McFarland for helpful discussions.

## AUTHOR CONTRIBUTIONS

MRW analyzed the data; MRW and DAB interpreted the results of experiments; MRW prepared the figures; MRW drafted the manuscript; MRW and DAB edited and revised the manuscript; MRW and DAB approved the final version of the manuscript.

---

**Figure A1.**
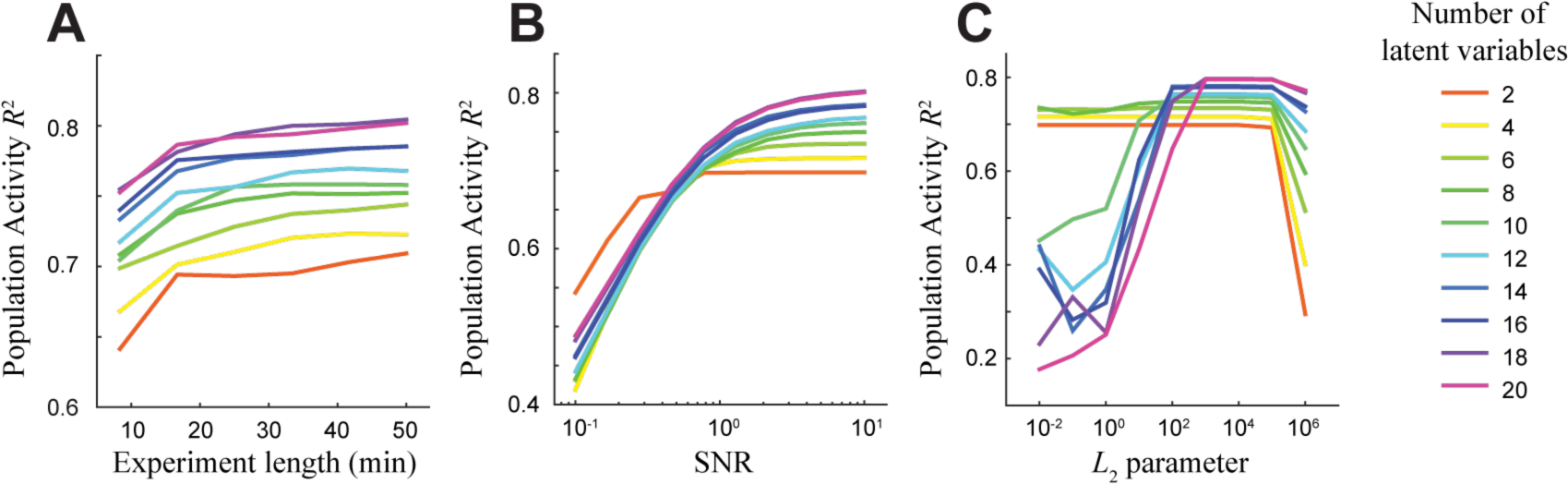
Sensitivity analysis of the autoencoder using simulated data. Datasets are generated as in Fig. 2 using varying numbers of latent variables. **(A-C)** An autoencoder is fit to each dataset using the correct number of latent variables. Plotted points represent the mean *R*^2^ value between the true and predicted population activity averaged over 20 such datasets; error bars are omitted for ease of interpretation. Plots show the result of varying: **A.** The amount of data used for fitting (using 10 Hz sampling rate); **B.** The signal-to-noise ratio of the data used for fitting (using 30 minutes of simulated data); **C.** The regularization parameter on the encoding and decoding weight matrices, which were constrained to be equal through weight-tying (again using 30 minutes of simulated data).

**Figure A2.**
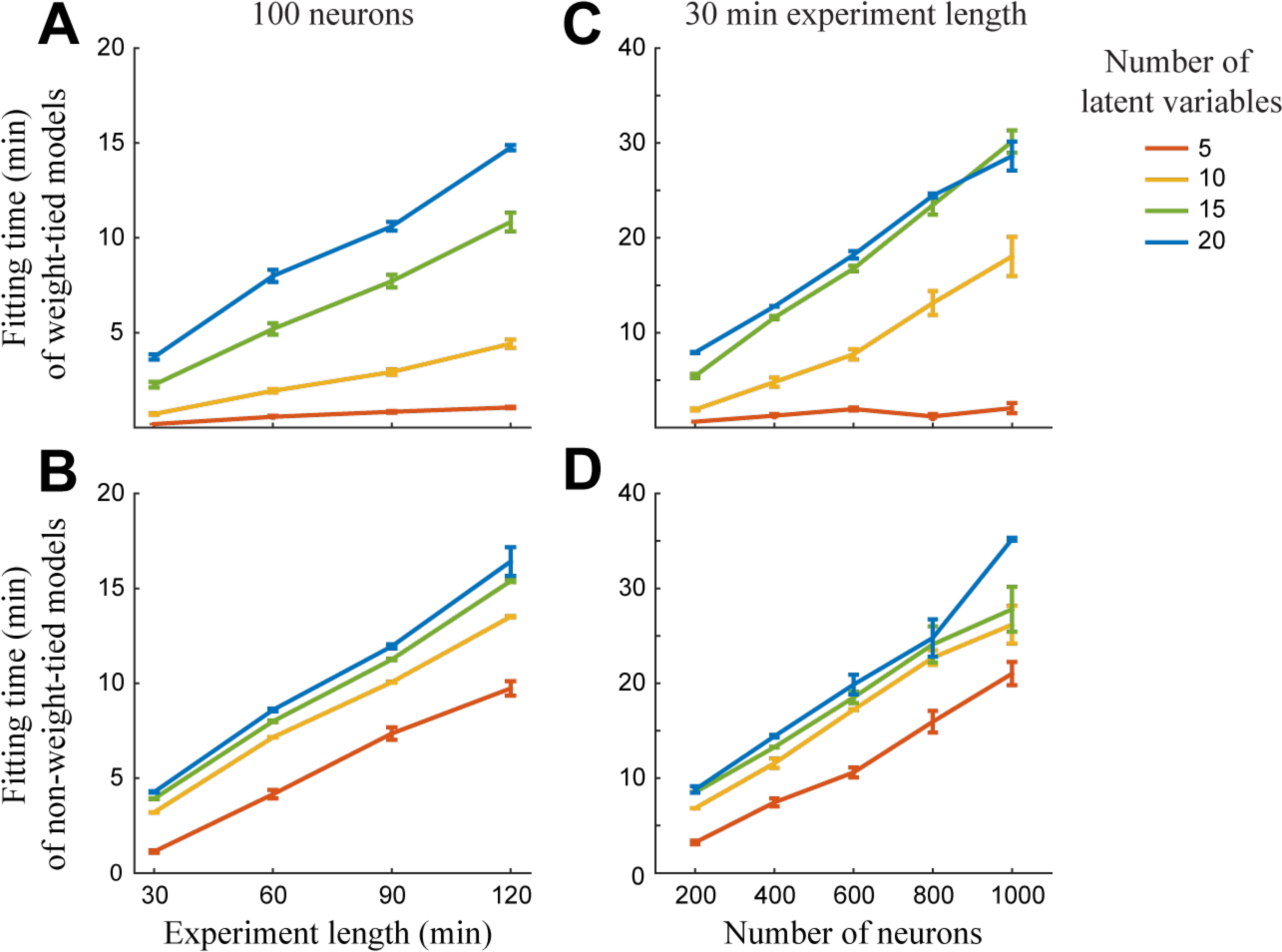
Linear scaling properties of the autoencoder. Plotted values are mean fitting times +/- standard error over 20 datasets. (**A-B**) Data is generated as in Fig. 2 with 100 neurons and varying recording lengths (with a 10 Hz sampling rate). Autoencoders are fit with and without weight-tying (A and B, respectively). The fitting time scales roughly linearly with the experiment time. (**C-D**) Data is generated as in Fig. 2 with a 30-minute experiment time and varying the number of neurons. Autoencoders are fit with and without weight-tying (C and D, respectively). The fitting time scales roughly linearly with the number of neurons. Comparing plots A and C (weight-tying) with plots B and D (no weight-tying) shows that while weight-tying approximately halves the number of estimated parameters, it leads to more than a two-fold speedup in fitting time with a small number of latent variables. As the number of latent variables increases this speedup advantage from weight-tying is lost. These results were obtained on a desktop machine running Ubuntu 14.04 LTS with 16 Intel^®^ Xeon^®^ E5-2670 processors and 126 GB of RAM; the MATLAB implementation of the autoencoder has not been optimized for this particular architecture.

**Figure A3.**
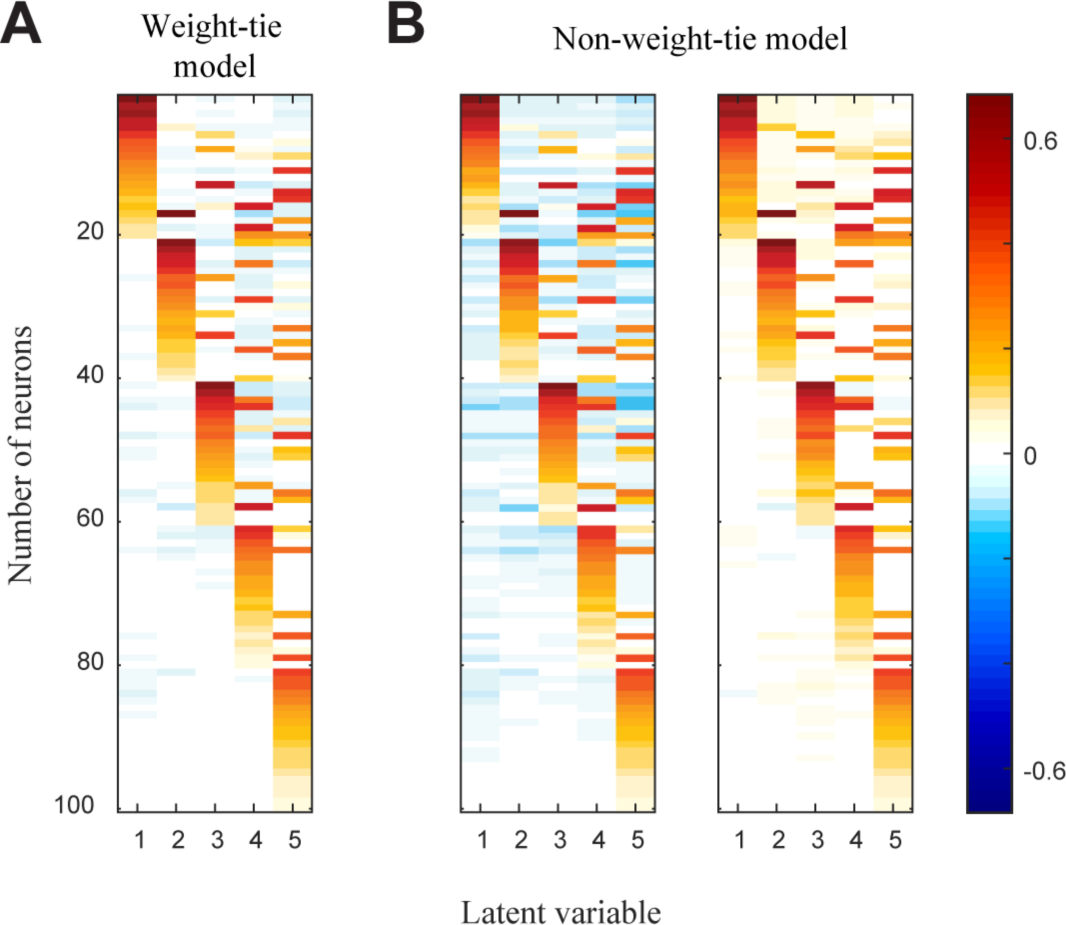
Effect of weight-tying using simulated data. We compared the effects of weight-tying the autoencoder on the resulting weight matrix by fitting models with and without weight-tying to the simulated data (Fig. 2). **A.** Weights learned by the autoencoder when encoding and decoding matrices are constrained to be the same. **B.** Encoding (*left*) and decoding (*right*) weights learned by the autoencoder without the weight-tying constraint, demonstrating a very similar pattern as the weight-tied solution in (A).

**Figure A4.**
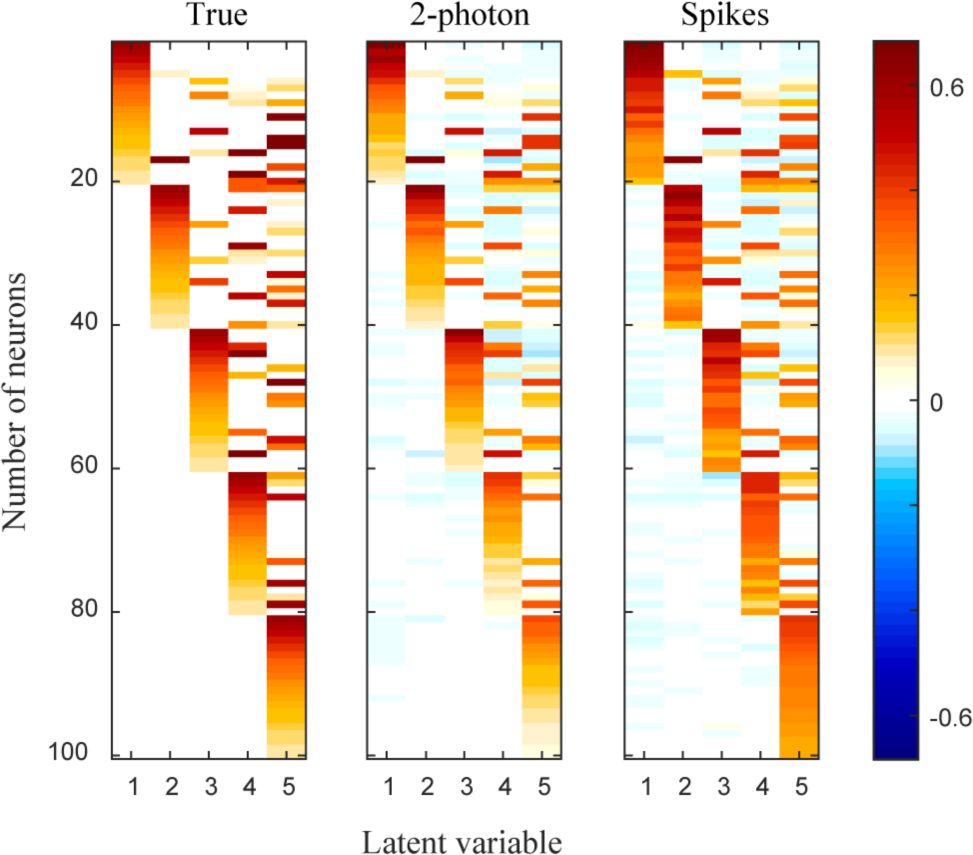
Using the RLVM for spiking data. *Right:*Coupling matrix used to generate synthetic data, as described in methods. *Middle*: The estimated coupling matrix when the autoencoder variant of the RLVM is fit to the simulated 2-photon data using a Gaussian noise loss function (mean square error). *Right:* The estimated coupling matrix when the autoencoder variant of the RLVM is fit to the simulated spiking data using a Poisson noise loss function (negative log-likelihood). The simulated data contained spikes binned at 100 ms resolution. The good agreement of both estimated coupling matrices with the true coupling matrix demonstrates that the RLVM can recover the same model parameters when fit using two different types of data. This result indicates that analyses similar to those presented in this paper can be used to equal effect on multielectrode data, without the need for data smoothing or averaging across trials (which are common preprocessing steps used with spiking data when attempting to use latent variable models not suited for discrete count data, such as PCA).

**Table A1.** Experimental selection. All experimental data used in Figs. 3-6 is from Peron et al. (2015) and is publicly available at http://dx.doi.org/10.6080/K0TB14TN. Data analysis was performed on subsets of the data that contained a large number of neurons simultaneously imaged over many trials (see Methods). The Analyzed Data column shows the amount of data retained from the Original Data column after removing neurons that had >50% missing values in their fluorescence traces or had an estimated SNR < 1, as well as removing trials that had missing values for any of the remaining fluorescence traces. The highlighted row corresponds to the dataset used for the analyses shown in Figs. 3-5.

|  | Original Data |  |  | Analyzed Data |  |
| --- | --- | --- | --- | --- | --- |
| Animal ID | Cell ID range | ROI count | Trial count | ROI count | Trial count |
| an229716 | 21000-23464 | 1395 | 157 | 116 | 135 |
|  | 15000-17352 | 1073 | 165 | 428 | 109 |
|  | 9000-11363 | 1006 | 160 | 176 | 135 |
| an229717 | 45000-47679 | 1695 | 173 | 698 | 146 |
|  | 21000-23694 | 1572 | 117 | 526 | 80 |
|  | 39000-41437 | 1313 | 136 | 496 | 93 |
| an229719 | 6000-8518 | 1496 | 161 | 106 | 115 |
|  | 9000-11423 | 1358 | 162 | 896 | 98 |
|  | 15000-17478 | 1332 | 158 | 484 | 113 |

## REFERENCES

Ahrens MB, Li JM, Orger MB, Robson DN, Schier AF, Engert F, Portugues R (2012) Brain-wide neuronal dynamics during motor adaptation in zebrafish. Nature 485: 471–477.

Amarasingham A, Geman S, Harrison MT (2015) Ambiguity and nonidentifiability in the statistical analysis of neural codes. Proc Natl Acad Sci USA 112: 6455–6460.

Archer EW, Koster U, Pillow JW, Macke JH (2014) Low-dimensional models of neural population activity in sensory cortical circuits. NIPS:343–351.

Arieli A, Sterkin A, Grinvald A, Aertsen A (1996) Dynamics of Ongoing Activity: Explanation of the Large Variability in Evoked Cortical Responses. Science 273: 1868–1871.

Bengio Y, Courville A, Vincent P (2013) Representation Learning: A Review and New Perspectives. IEEE Transactions on Pattern Analysis and Machine Intelligence 35: 1798–1828.

Bishop CM (2006) Pattern recognition. Machine Learning.

Boulard H, Kamp Y (1989) Autoassociative memory by multilayer perceptron and singular values decomposition. Biological Cybernetics 59: 291–294.

Churchland MM et al. (2010) Stimulus onset quenches neural variability: a widespread cortical phenomenon. Nature Neuroscience 13: 369–378.

Churchland MM, Cunningham JP, Kaufman MT, Foster JD, Nuyujukian P, Ryu SI, Shenoy KV (2012) Neural population dynamics during reaching. Nature 487: 51–56.

Cohen MR, Kohn A (2011) Measuring and interpreting neuronal correlations. Nature Neuroscience 14: 811–819.

Cui Y, Liu LD, McFarland JM, Pack CC, Butts DA (2016) Inferring Cortical Variability from Local Field Potentials. J Neurosci 36: 4121–4135.

Cunningham JP, Yu BM (2014) Dimensionality reduction for large-scale neural recordings. Nature Neuroscience 17: 1500–1509.

De Meo R, Murray MM, Clarke S, Matusz PJ (2015) Top-down control and early multisensory processes: chicken vs. egg. Front Integr Neurosci 9: 1–6.

Doiron B, Litwin-Kumar A, Rosenbaum R, Ocker GK, Josić K (2016) The mechanics of state-dependent neural correlations. Nature Neuroscience 19: 383–393.

Freeman J, Vladimirov N, Kawashima T, Mu Y, Sofroniew NJ, Bennett DV, Rosen J, Yang C-T, Looger LL, Ahrens MB (2014) Mapping brain activity at scale with cluster computing. Nat Meth 11: 941–950.

Ghazanfar A, Schroeder C (2006) Is neocortex essentially multisensory? Trends in Cognitive Sciences 10: 278–285.

Goris RLT, Movshon JA, Simoncelli EP (2014) Partitioning neuronal variability. Nature Neuroscience 17: 858–865.

Haggerty DC, Ji D (2015) Activities of visual cortical and hippocampal neurons co-fluctuate in freely moving rats during spatial behavior. eLife Sciences 4:e08902.

Hara K, Saito D, Shouno H (2015) Analysis of function of rectified linear unit used in deep learning. 2015 International Joint Conference on Neural Networks (IJCNN):1–8.

Harris KD, Thiele A (2011) Cortical state and attention. Nature Reviews Neuroscience 12: 509–523.

Hinton GE (2012) A Practical Guide to Training Restricted Boltzmann Machines. In: Neural Networks: Tricks of the Trade, pp 599–619 Lecture Notes in Computer Science. Berlin, Heidelberg: Springer Berlin Heidelberg.

Japkowicz N, Hanson SJ, Gluck MA (2000) Nonlinear Autoassociation Is Not Equivalent to PCA. Neural Computation 12: 531–545.

Kato S, Kaplan HS, Schrödel T, Skora S, Lindsay TH, Yemini E, Lockery S, Zimmer M (2015) Global Brain Dynamics Embed the Motor Command Sequence of Caenorhabditis elegans. Cell 163: 656–669.

Koster U, Sohl-Dickstein J, Gray CM, Olshausen BA (2014) Modeling Higher-Order Correlations within Cortical Microcolumns Macke JH, ed. PLoS Comput Biol 10:e1003684.

Kulkarni JE, Paninski L (2015) Common-input models for multiple neural spike-train data. Network: Computation in Neural Systems 18: 375–407.

Lakshmanan KC, Sadtler PT, Tyler-Kabara EC, Batista AP, Yu BM (2015) Extracting Low-Dimensional Latent Structure from Time Series in the Presence of Delays. Neural Computation 27: 1825–1856.

Lee DD, Seung HS (1999) Learning the parts of objects by non-negative matrix factorization. Nature 401: 788–791.

Lin I-C, Okun M, Carandini M, Harris KD (2015) The Nature of Shared Cortical Variability. Neuron 87: 644–656.

Macke JH, Buesing L, Cunningham JP, Yu BM, Shenoy KV, Sahani M (2011) Empirical models of spiking in neural populations. NIPS:1350–1358.

Marguet SL, Harris KD (2011) State-Dependent Representation of Amplitude-Modulated Noise Stimuli in Rat Auditory Cortex. J Neurosci 31: 6414–6420.

McFarland JM, Cui Y, Butts DA (2013) Inferring Nonlinear Neuronal Computation Based on Physiologically Plausible Inputs Bethge M, ed. PLoS Comput Biol 9:e1003143.

Niell CM, Stryker MP (2010) Modulation of Visual Responses by Behavioral State in Mouse Visual Cortex. Neuron 65: 472–479.

Ohiorhenuan IE, Mechler F, Purpura KP, Schmid AM, Hu Q, Victor JD (2010) Sparse coding and high-order correlations in fine-scale cortical networks. Nature 466: 617–621.

Okun M, Steinmetz NA, Cossell L, Iacaruso MF, Ko H, Barthó P, Moore T, Hofer SB, Mrsic-Flogel TD, Carandini M, Harris KD (2015) Diverse coupling of neurons to populations in sensory cortex. Nature 521: 511–515.

Otazu GH, Tai L-H, Yang Y, Zador AM (2009) Engaging in an auditory task suppresses responses in auditory cortex. Nature Neuroscience 12: 646–654.

Pachitariu M, Lyamzin DR, Sahani M, Lesica NA (2015) State-Dependent Population Coding in Primary Auditory Cortex. J Neurosci 35: 2058–2073.

Paninski L (2004) Maximum likelihood estimation of cascade point-process neural encoding models. Network: Computation in Neural Systems 15: 243–262.

Paninski L, Ahmadian Y, Ferreira DG, Koyama S, Rahnama Rad K, Vidne M, Vogelstein J, Wu W (2009) A new look at state-space models for neural data. J Comput Neurosci 29: 107–126.

Peron SP, Freeman J, Iyer V, Guo C, Svoboda K (2015) A Cellular Resolution Map of Barrel Cortex Activity during Tactile Behavior. Neuron 86: 783–799.

Pfau D, Pnevmatikakis EA, Paninski L (2013) Robust learning of low-dimensional dynamics from large neural ensembles. NIPS:2391–2399.

Pillow JW, Shlens J, Paninski L, Sher A, Litke AM, Chichilnisky EJ, Simoncelli EP (2008) Spatio-temporal correlations and visual signalling in a complete neuronal population. Nature 454: 995–999.

Rabinowitz NC, Goris RL, Cohen M, Simoncelli EP (2015) Attention stabilizes the shared gain of V4 populations. eLife Sciences 4:e08998.

Rasch MJ, Gretton A, Murayama Y, Maass W, Logothetis NK (2008) Inferring Spike Trains From Local Field Potentials. Journal of Neurophysiology 99: 1461–1476.

Schneidman E, Berry MJ, Segev R, Bialek W (2006) Weak pairwise correlations imply strongly correlated network states in a neural population. Nature 440: 1007–1012.

Schölvinck ML, Saleem AB, Benucci A, Harris KD, Carandini M (2015) Cortical State Determines Global Variability and Correlations in Visual Cortex. J Neurosci 35: 170–178.

Schultz W, Carelli RM, Wightman RM (2015) Phasic dopamine signals: from subjective reward value to formal economic utility. Current Opinion in Behavioral Sciences 5: 147–154.

Semedo J, Zandvakili A, Kohn A, Machens CK, Yu BM (2014) Extracting Latent Structure From Multiple Interacting Neural Populations. NIPS:2942–2950.

Shuler MG (2006) Reward Timing in the Primary Visual Cortex. Science 311: 1606–1609.

Smith AC, Brown EN (2003) Estimating a State-Space Model from Point Process Observations. Neural Computation 15: 965–991.

Stopfer M, Jayaraman V, Laurent G (2003) Intensity versus Identity Coding in an Olfactory System. Neuron 39: 991–1004.

Vidne M, Ahmadian Y, Shlens J, Pillow JW, Kulkarni J, Litke AM, Chichilnisky EJ, Simoncelli E, Paninski L (2011) Modeling the impact of common noise inputs on the network activity of retinal ganglion cells. J Comput Neurosci 33: 97–121.

Vinck M, Batista-Brito R, Knoblich U, Cardin JA (2015) Arousal and Locomotion Make Distinct Contributions to Cortical Activity Patterns and Visual Encoding. Neuron 86: 740–754.

Yu BM, Cunningham JP, Santhanam G, Ryu SI, Shenoy KV, Sahani M (2009) Gaussian-Process Factor Analysis for Low-Dimensional Single-Trial Analysis of Neural Population Activity. Journal of Neurophysiology 102: 614–635.

